# Cell membranes resist flow

**DOI:** 10.1101/290643

**Authors:** Zheng Shi, Zachary T. Graber, Tobias Baumgart, Howard A. Stone, Adam E. Cohen

## Abstract

The fluid-mosaic model posits a liquid-like plasma membrane, which can flow in response to tension gradients. It is widely assumed that membrane flow transmits local changes in membrane tension across the cell in milliseconds. This conjectured signaling mechanism has been invoked to explain how cells coordinate changes in shape, motility, and vesicle fusion, but the underlying propagation has never been observed. Here we show that propagation of membrane tension occurs quickly in cell-attached blebs, but is largely suppressed in intact cells. The failure of tension to propagate in cells is explained by a fluid dynamical model that incorporates the flow resistance from cytoskeleton-bound transmembrane proteins. In primary endothelial cells, local increases in membrane tension lead only to local activation of mechanosensitive ion channels and to local vesicle fusion. Thus membrane tension is not a mediator of long-range intra-cellular signaling, but local variations in tension mediate distinct processes in sub-cellular domains.

## INTRODUCTION

Membrane tension affects cell migration (Gauthier et al., 2011; Houk et al., 2012; Keren et al., 2008), vesicle fusion and recycling (Boulant et al., 2011; Gauthier et al., 2011; Maritzen and Haucke, 2017; Shillcock and Lipowsky, 2005), the cell cycle (Stewart et al., 2011), cell signaling (Basu et al., 2016; Groves and Kuriyan, 2010; Romer et al., 2007), and mechanosensation (Phillips et al., 2009; Ranade et al., 2015). In artificial lipid bilayers, changes in membrane tension propagate across a cell-sized region in milliseconds (Fig. S1) (Shi and Baumgart, 2015). Fluorescently tagged transmembrane proteins typically diffuse freely in both artificial bilayers and in intact cells, albeit with a 10-100 fold lower diffusion coefficient in cells (Kusumi et al., 2005). Together these results, each consistent with the fluid mosaic model (Singer and Nicolson, 1972), led to the widespread belief that two-dimensional (2D) flow of lipids in cells mediates rapid intracellular equilibration of membrane tension (Keren et al., 2008; Kozlov and Mogilner, 2007), providing a long-range signaling mechanism analogous to the rapid propagation of electrical signals in neurons (Keren, 2011).

Intact cell membranes contain many features not found in artificial lipid bilayers. In eukaryotic cells, approximately half of the transmembrane proteins, corresponding to ∼10 - 20% of total membrane area (Bussell et al., 1995; Zakharova et al., 1995), are bound to the underlying cortex and therefore are effectively immobile on timescales of minutes to hours (Bussell et al., 1995; Groves and Kuriyan, 2010; Sadegh et al., 2017). Are these obstacles a minor perturbation to lipid flow, or do they qualitatively change the dynamics? Aqueous solutions with ∼10% immobile protein, such as collagen gels, behave as bulk solids, not liquids, yet still permit lateral diffusion of small molecules and proteins. Thus it is plausible that cell membranes, too, could exist in a state that behaves as a 2D fluid on the nanoscale but that is closer to a semi-solid gel on the cellular scale. The 2D-gel hypothesis is incompatible with the many conjectures in the literature that rapid propagation of membrane tension can mediate long-range intracellular signaling (Diz-Muñoz et al., 2013; Gauthier et al., 2012; Gauthier et al., 2011; Houk et al., 2012; Keren et al., 2008; Keren, 2011; Morris and Homann, 2001; Mueller et al., 2017; Pontes et al., 2017; Watanabe et al., 2013).

## RESULTS

### Membrane tension propagates in membrane blebs but not in cell membranes

Working with HeLa cells at 37 °C, we pulled short membrane tethers as a means of simultaneously perturbing and measuring local membrane tension (Fig. 1A, 1D). Tether diameter and local membrane tension are inversely related, coupled via the membrane’s finite bending stiffness (*Supplementary Discussion*) (Derényi et al., 2002; Pontes et al., 2017). Tether diameters were too small to resolve optically, so we used fluorescence of a membrane-bound tag (GPI-eGFP) to estimate tether diameter. Under wide-field fluorescence excitation, flare from the cell body prevented accurate quantification of the fluorescence from the far dimmer tether. We used a custom micromirror-based patterned illumination system to restrict fluorescence excitation to the tethers, leading to high-contrast images of individual tethers. By calibrating tether fluorescence against the fluorescence of a patch of cell membrane with known area, we determined the tether diameter. We used simultaneous fluorescence and optical trap force measurements to calibrate the relationship between tether diameter and tension (Fig. S2).

**Fig. 1.**
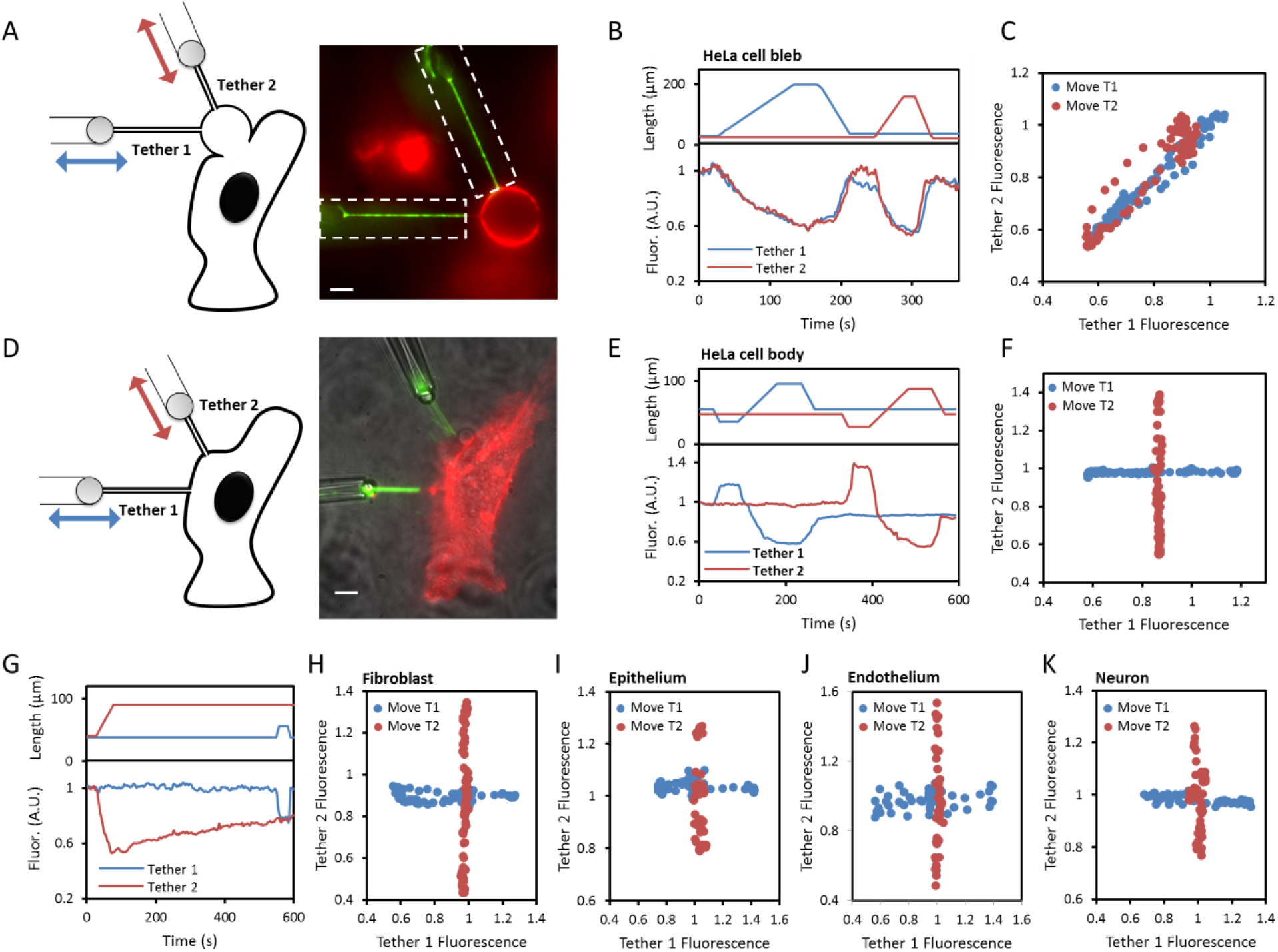
Propagation of membrane tension in cells. **A,D)** Schematic (left) and fluorescence image (right) showing a pair of tethers pulled from **(A)** a cell-attached bleb or **(D)** the cell body of a HeLa cell expressing GPI-eGFP. Green: fluorescence under patterned illumination (restricted to dashed boxes). Red: fluorescence under wide-field illumination. In **(D)** a transmitted light image (grey) is combined with the fluorescence images. Scale bars 5 μm. **B,E)** The two tethers were stretched sequentially (top) and the fluorescence of each tether was monitored (bottom). **C,F)** Relation between the intensities of the two tethers when either the first or second tether was stretched. **G)** Test for slow coupling between tethers in a HeLa cell. A change in length of tether 2 did not affect fluorescence of tether 1 within a 500 s measurement window. **H-K)** Repetition of the experiment in **(D-F)** in **H)** NIH 3T3 fibroblasts, **I)** MDCK epithelial cells, **J)** mouse brain endothelial cells, and **K)** rat hippocampal neurons. T1: Tether 1, T2: Tether 2.

Two membrane tethers were then simultaneously pulled from nearby locations on a single cell (typically 5-15 µm apart) and fluorescence from each was excited with micromirror-patterned illumination (Fig. 1A, 1D). Each tether was successively stretched and relaxed, while the fluorescence of both tethers was monitored to measure local tension. In cell-attached membrane blebs, we observed tight coupling of the tension in the two tethers (Fig. 1A - C). Stretching of either tether led to a decrease in the fluorescence of both tethers, with the response of the unstretched tether lagging by < 1 s. Measurements on 10 pairs of tethers pulled from different blebs all showed strongly coupled fluorescence changes. Thus tension rapidly equilibrated across blebs, consistent with observations in artificial lipid vesicles (e.g. Fig. S1). In intact cells, in contrast, we failed to observe any coupling between the tethers (Fig. 1D - F). Measurements lasted up to 500 s, and attachment points were as close as 5 µm (Fig. 1G). We tested HeLa cells (Fig. 1E, 1F, *n* = 30 cells), NIH 3T3 fibroblasts (Fig. 1H, *n* = 10 cells), MDCK epithelial cells (Fig. 1I, *n* = 5 cells), mouse brain endothelial cells (Fig. 1J, *n* = 5 cells), and proximal dendrites of rat hippocampal neurons (Fig. 1K, *n* = 5 neurons) and did not observe tension propagation in any of these cell types.

The failure to observe propagation of membrane tension in cells might be explained by rapid assembly of cytoskeletal barriers that isolated the tether from the rest of the cell. To test for such barriers, we performed fluorescence recovery after photobleaching (FRAP) experiments in the tethers immediately adjacent to the attachment point (Lippincott-Schwartz et al., 2003). The fluorescence recovery profile quantitatively matched simulations of free diffusion between the cell and tether, ruling out local cytoskeletal isolation of the tether (Fig. S3).

### Hydrodynamic model of membrane flow

In two-dimensional flows, an immobile obstacle creates a logarithmically diverging long-range perturbation to the flow field, a phenomenon sometimes called the Stokes paradox. We hypothesized that cytoskeleton-bound transmembrane proteins might significantly impede the membrane flow required to propagate tension changes in cells (Fig. 2A). Over length scales large compared to the inter-obstacle spacing, the poroelastic equations governing lipid flow lead to a diffusion-like equation for propagation of membrane tension, with tension diffusion coefficient *D*_*σ*_ = *E*_*m*_*k*/*η*, where *E*_m_ is the effective area expansion modulus of the membrane, *η* is the two-dimensional membrane viscosity, and *k* is the Darcy permeability of the array of obstacles (*Supplementary Discussion*). The diffusion coefficient for the spread of membrane tension represents the balance of viscous and elastic forces in the membrane (Fig. 2B), and is physically distinct from the diffusion coefficients that govern motion of tracer molecules within the lipid bilayer.

**Fig. 2.**
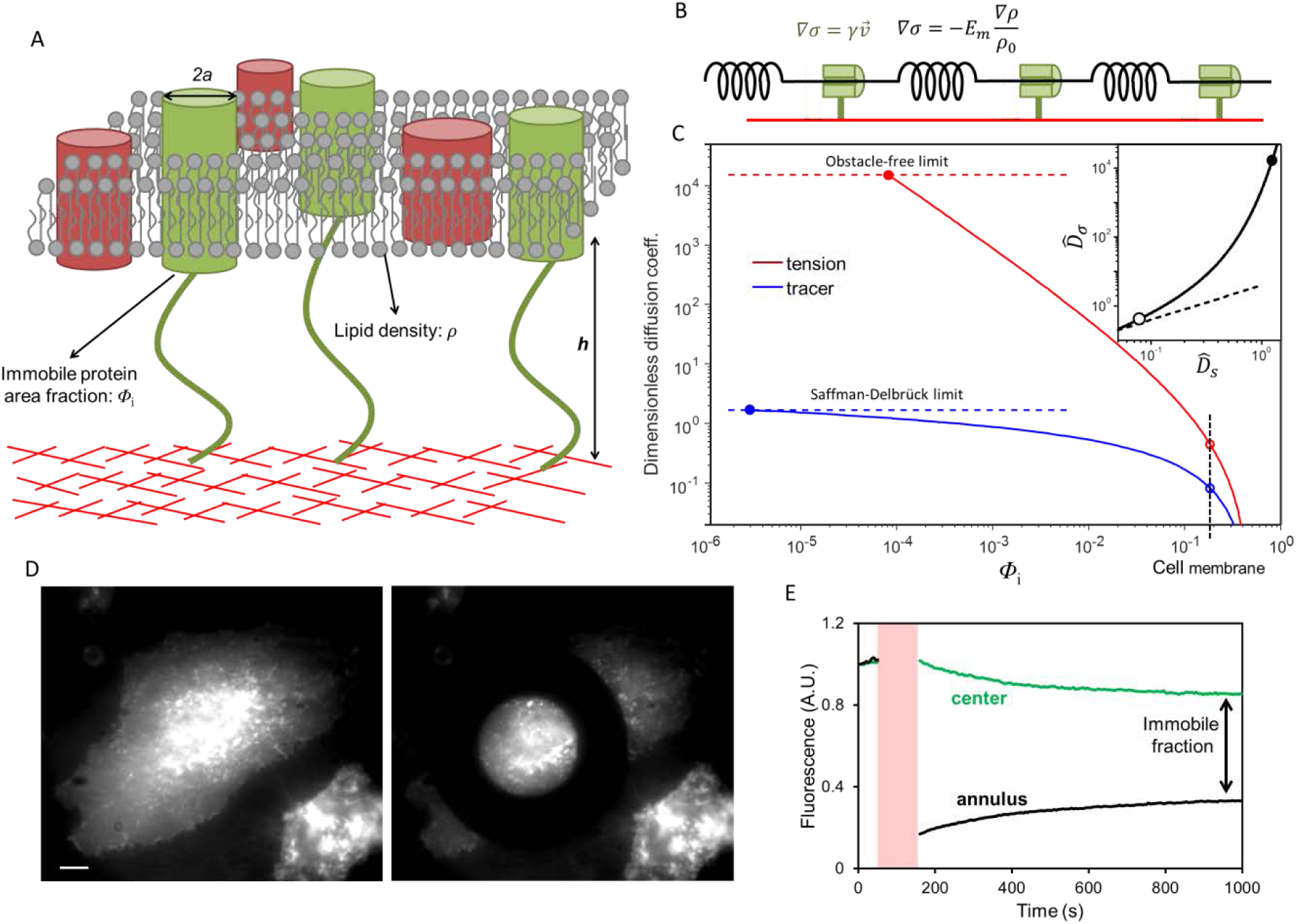
Hydrodynamic model of membrane flow past immobile obstacles. **A)** Illustration of the cell plasma membrane with some transmembrane proteins bound to the underlying cortex. **B)** Simple viscoelastic model of the cell membrane. Springs represent the elastic response of the membrane to stretch, and dampers represent the viscous drag from immobile transmembrane proteins. **C)** Dependence of diffusion coefficients for membrane tension (red) and molecular tracers (blue) on the area fraction *Φ*_i_ of immobile proteins. This plot shows the model’s predictions for the dimensionless diffusion coefficients, 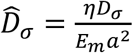 for tension and 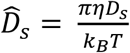 for tracers. The upper limit on tension diffusion is set by the hydrodynamic drag between plasma membrane and cytoskeleton cortex in the absence of obstacles. The upper limit on tracer diffusion is set by the Saffman–Delbrück model (*Supplementary Discussion*). Open circles: diffusion coefficients in intact cell membranes. Inset: Relation between dimensionless diffusion coefficients of membrane tension and molecular tracers (solid line). The dashed line shows a linear relation. Closed circles: obstacle-free membrane. Open circles: *Φ*_i_ = 0.18. **D)** Fluorescence image showing a HeLa cell in which transmembrane proteins have been labeled non-specifically with Alexa488-NHS before (left) and after (right) bleaching with a donut shape laser spot. Scale bar, 10 μm. **E)** Fluorescence intensity profile of the bleached ring (black) and non-bleached central (green) regions. The photobleaching epoch is shaded red.

The Darcy permeability, *k*, scales with obstacle radius, *a* and area fraction of the obstacles, *Φ*_i_, as *k* = *a*^2^*f*(*Φ*_i_), where *f*(*Φ*_i_) is a dimensionless function which varies steeply at small *Φ*_i_ (Fig. S4) (Bussell et al., 1995; Howells, 1974). Bussell *et al.* (Bussell et al., 1995) showed that one can estimate *Φ*_i_ from the diffusion of transmembrane tracer molecules. We compared the diffusion coefficients, *D*_*S*_, of tracer molecules on an intact cell versus on cytoskeleton-free membrane tethers (Fig. 2C and Fig. S5). For a transmembrane dopamine receptor fused to eGFP (DRD2-eGFP), FRAP measurements yielded diffusion coefficients on the cell 21 ± 4 fold lower than on the tether (*D*_S_^cell^ = 0.037 ± 0.005 μm^2^/s, *D*_S_^tether^ = 0.76 ± 0.08 μm^2^/s, mean ± s.e.m., *n* = 10 pairs of tethers and cells). We explored a variety of other tracers to control for possible molecularly specific interactions with cytoskeletal components and obtained similar results (Table S1), consistent with literature (Kusumi et al., 2005). We used the Saffman– Delbrück model (Saffman and Delbrück, 1975) to fit the diffusion on the cytoskeleton-free tethers, and the Bussel model (Bussell et al., 1995) to fit the diffusion on the cell body. The pair of fits yielded a membrane viscosity *η* = (3.0 ± 0.4) × 10^-3^ pN·s/μm and an area fraction of immobile obstacles *Φ*_*i*_ = 0.18 ± 0.03 (Fig. 2C), consistent with literature results (Bussell et al., 1995; Kusumi et al., 2005).

We performed additional FRAP experiments to make an independent estimate of *Φ*_*i*_ in HeLa cells. Transmembrane proteins were labeled nonspecifically with a broadly reactive cell-impermeant dye, and then photobleached in a sub-cellular region (Fig. 2D). Mobile proteins thereafter diffused back into the bleached region, while immobile proteins did not. The degree of partial fluorescence recovery at long time (15 min) showed that 54 ± 5% (mean ± s.e.m., *n* = 5 cells) of all labeled transmembrane proteins were immobile (Fig. 2E, Fig. S6). When combined with literature estimates that ∼25% of membrane area is occupied by transmembrane proteins (Dupuy and Engelman, 2008; Zakharova et al., 1995), these results are broadly consistent with our estimate of *Φ*_i_ = 0.1 – 0.2 based on molecular diffusion measurements.

Combining the estimates of membrane viscosity, *η* = (3.0 ± 0.4) × 10^-3^ pN·s/μm, and obstacle area fraction (*Φ*_*i*_ = 0.18 ± 0.03) with reasonable values of the obstacle radius (*a* ∼ 2 nm) (Bussell et al., 1995) and effective membrane area expansion modulus (*E*_*m*_ = 40 pN/μm) (Hochmuth, 2000) yields a tension diffusion coefficient: *D*_σ_ = 0.024 ± 0.005 μm^2^/s. Tension therefore requires tens of minutes to equilibrate over cellular length scales (∼10 µm). These experiments additionally yielded an estimate of an effective drag coefficient for membrane flow relative to the cytoskeleton, γ = *η/k*. We found γ = (1700 ± 300) pN·s/μm^3^. Microrheometry measurement of cell plasma membrane with magnetic particles yielded a similar drag coefficient, γ ≈ 2000 pN·s/μm^3^ (Bausch et al., 1998).

The hydrodynamic model establishes that the tension diffusion coefficient *D*_σ_ is far more sensitive to obstacles than is the tracer diffusion coefficient, *D*_*S*_. An obstacle density that decreases tracer diffusion 10-fold from the free-membrane limit, decreases diffusion of tension 10^4^-fold (Fig. 2C and *Supplementary Discussion*). Obstacles at densities that modestly suppress tracer diffusion will almost completely block lipid flow, causing the membrane to appear rheologically like a gel.

The model also predicts the rate at which lipids flow into a tether after a ramp increase in tether length. Using the experimentally determined tension diffusion coefficient (D_σ_ = 0.024 μm^2^/s), we simulated this process, accounting for the gradual changes in tether tension and radius as lipid flowed into the tether. These simulations, which had no adjustable parameters, quantitatively matched the measurements of the time-dependent tether radius (Fig. S7). The simulations showed that the perturbation to membrane tension remained localized near the tether. At a radius of 5 µm from the tether, the maximum increase in tension was less than 3% of the maximum increase at the tether attachment point. Thus local perturbations to membrane tension remained localized.

### Localized activation of mechanosensitive ion channels (MSCs) and vesicle fusion

To test whether membrane tension is a local or global regulator of membrane signaling, we examined the effect of local perturbations to tension on the activation of mechanosensitive ion channels. We pulled tethers in endogenously mechanosensitive MDCK cells (Gudipaty et al., 2017), and performed simultaneous dual-color imaging of tether fluorescence (via GPI-eGFP) and intracellular Ca^2+^ (via R-CaMP2, Fig. 3A-C). These experiments revealed that MSCs in MDCK cells activated at a membrane tension ∼10-fold higher than the resting tension (Fig. S8). We then switched to using GCaMP6f to improve Ca^2+^ sensitivity. In 18 out of 27 trials (15 out of 21 cells) tether pulling triggered Ca^2+^ influx (Fig. 3D). In all tether pulling experiments the Ca^2+^ influx, if detected, occurred at the tether attachment to within our ability to resolve these two sites (Fig. 3F). We never observed tether-induced Ca^2+^ influx in any regions of the cell distal from the tether attachment, even at the slowest pulling speeds tested (1 µm/s), establishing that membrane tension acted locally, not globally, to gate endogenous MSCs.

**Fig. 3.**
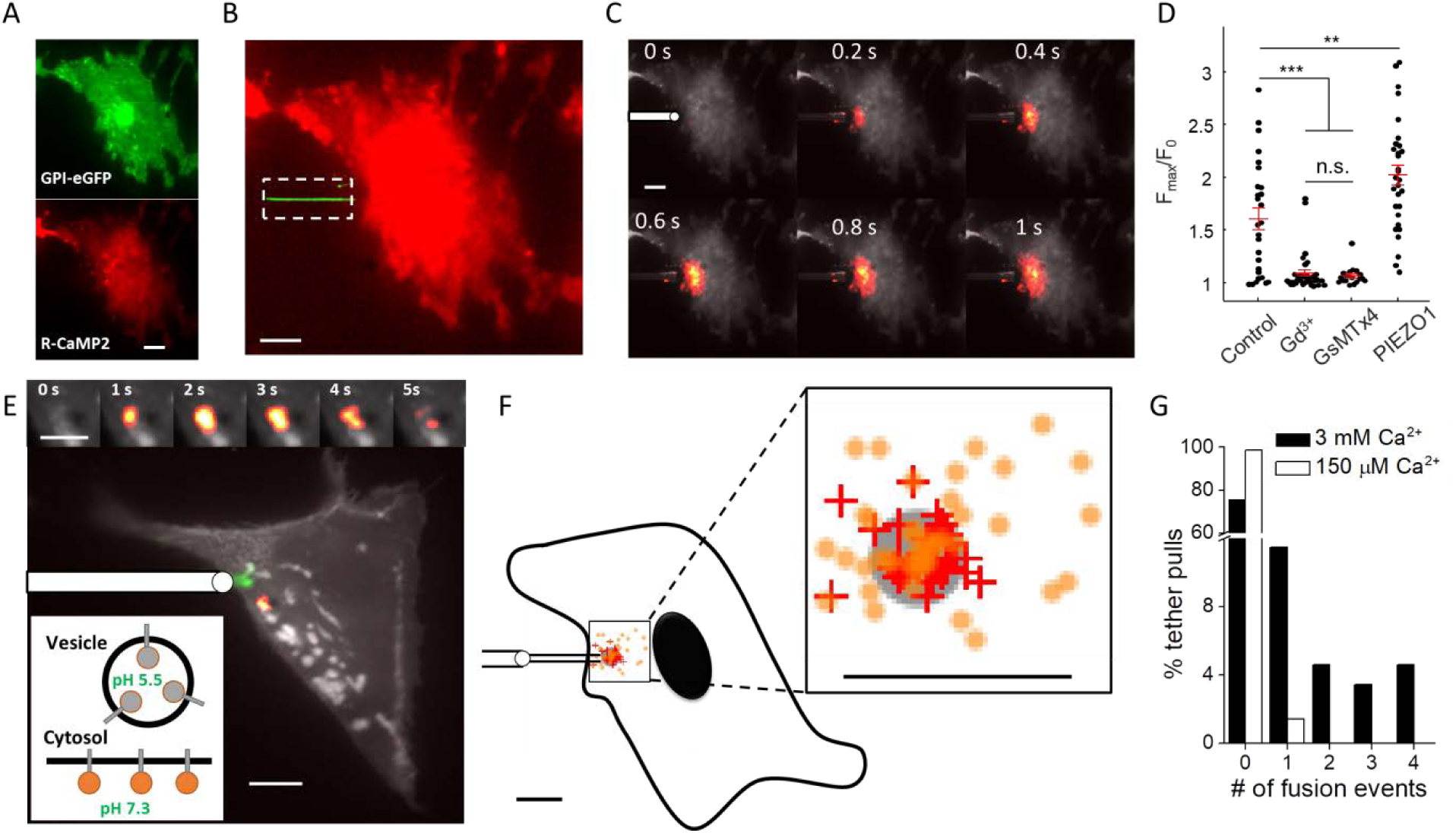
Membrane tension mediates local activation of mechanosensitive ion channels and local vesicle fusion in MDCK cells. **A)** MDCK cell co-expressing GPI-eGFP (green) and R-CaMP2 (red). **B)** Composite fluorescence image of tether (green) and R-CaMP2 (red). Fluorescence excitation of eGFP was confined to the tether (dashed box). **C)** Localized Ca^2+^ influx triggered by tether stretch. Images are composites of mean fluorescence (grey) and changes in fluorescence (heat map). Tether pulling pipette shown schematically at 0 s. **D)** Blockers of MSCs, GdCl_3_ (500 µM) or GsMTx4 (8 µM), suppressed Ca^2+^ influx during tether pulling. Over-expression of PIEZO1-mCherry increased Ca^2+^ influx during tether pulling (*n* = 27 cells in control extracellular buffer, *n* = 36 with GdCl_3_, *n* = 18 with GsMTx4, *n* = 31 with PIEZO1 over-expression, ** *p*<0.01, *** *p* < 10^-3^, n.s.: *p* > 0.5, Student’s t-test). Data points represent maximal fractional increase in fluorescence of Ca^2+^ reporter. Red lines: mean. Error bars: s.e.m. **E)** Composite fluorescence image of mean fluorescence (grey), changes in fluorescence after tether pull (heat map) and tether location (green). Tether pulling pipette shown schematically. Upper inset: close-up view of the vesicle fusion events triggered by tether stretch. Lower inset: membrane-tethered mOrange2-TM reported vesicle fusion via pH-mediated changes in fluorescence. **F)** Distribution of Ca^2+^ influx (+) and vesicle fusion (o) sites relative to the tether attachment point (grey circle). Each mark represents one event (33 Ca^2+^ influx events from 25 cells; 43 vesicle fusion events from 21 cells). Average distance between Ca^2+^ initiation and tether attachment was 1.7 ± 0.2 μm (mean ± s.e.m), smaller than the localization uncertainty (3 μm). Average distance between vesicle fusion site and tether attachment was 3.5 ± 0.4 μm (mean ± s.e.m), much smaller than the the null hypothesis of uniform fusion throughout the cell (27 ± 2 μm). The outline of the cell is a schematic to illustrate size. **G)** In control extracellular medium (3 mM Ca^2+^) tether pulling triggered fusion of one or more vesicles in 21 out of 87 trials (black). In low [Ca^2+^] buffer (150 µM Ca^2+^ buffered by EGTA) tether pulling triggered fusion of only one vesicle in 71 trials (white). Scale bars in all panels 10 μm, except 5 μm for the upper inset in (E).

The Ca^2+^ influx was largely blocked by Gd^3+^ (2 Ca^2+^ influx events in 36 tether pulls, Fig. 3D) confirming that the influx was through stretch-activated mechanosensitive ion channels (Hua et al., 2010). The peptide toxin GsMTx4 also blocked the tether-induced Ca^2+^ influx (1 Ca^2+^ influx event in 18 tether pulls, Fig. 3D). This toxin blocks PIEZO1 but not other MSCs such as TREK-1 (Bae et al., 2011), suggesting that PIEZO1 likely mediates localized tension sensing in MDCK cells. Overexpression of PIEZO1-mCherry in MDCK cells led to increased but still localized Ca^2+^ influx during tether pulling (Fig. 3D, Fig. S9), confirming that PIEZO1 reports local, not global, membrane tension (Saotome et al., 2017). Sequential tether pulling from different locations of the same cell led to local Ca^2+^ influx at each pulling location but not at the previously pulled site, further demonstrating sub-cellular compartmentalization of mechanosensation (Fig. S9C).

Increases in membrane tension have been reported to facilitate vesicle release (Gauthier et al., 2011; Shillcock and Lipowsky, 2005). We next tested whether this effect was local or global. We expressed in MDCK cells membrane-tethered mOrange2 (mOrange2-TM), targeted to the inside of vesicles and to the extracellular face of the plasma membrane (*Materials and Methods*). This pH-sensitive reporter (pK_a_ 6.5 (Shaner et al., 2008)) was dark in the acidic lumen of vesicles and became fluorescent upon vesicle fusion to the plasma membrane (Fig. 3E). Addition of the Ca^2+^ ionophore ionomycin (5 μM) led to Ca^2+^ influx and vesicle fusion as reported by the dye FM 4-64, confirming that ionomycin triggered vesicle release (Fig. S10). In MDCK cells expressing mOrange2-TM, ionomycin led to a cell-wide appearance of bright fluorescent puncta, confirming the ability of mOrange2-TM to report vesicle fusion (Fig. S10). We then pulled tethers (from fresh cells expressing mOrange2-TM) and mapped the distribution of ensuing vesicle fusion events (Fig. 3E,F). We compared to the distribution anticipated from the null hypothesis of uniform fusion throughout the cell (*Materials and Methods*). The tension-induced events were significantly clustered around the tethers (Fig. 3F, mean distance 3.5 ± 0.4 μm, vs. 27 ± 2 μm for null hypothesis, mean ± s.e.m, *n* = 43 fusion events from 21 cells).

The vesicle fusion events were more broadly distributed around the tether attachment points than were the Ca^2+^ influx sites (*p* = 0.001), leading us to hypothesize that the vesicle fusion might be predominantly mediated by Ca^2+^ influx at the tether attachment and then Ca^2+^ diffusion over a larger, but still sub-cellular, region. Indeed, buffering extracellular Ca^2+^ concentration to 150 μM with EGTA largely eliminated tension-induced vesicle fusion (Fig. 3G, only 1/71 pulls induced fusion), establishing that the local vesicle fusion was mediated by local influx of Ca^2+^ through MSCs.

Endothelial cells respond to changes in shear flow *in vivo* (Geiger et al., 1992; Li et al., 2014; Schwarz et al., 1992) via activation of PIEZO1 (Guo and MacKinnon, 2017). We thus asked whether tension-induced activation of mechanosensitive ion channels in primary mouse brain endothelial cells (mBECs) was local or global. As in the MDCK cells, tether pulling led to local influx of Ca^2+^ and local vesicle fusion (Fig. 4A-C). Tethers are a non-physiological perturbation, so we then tested the effect of localized shear flow on Ca^2+^ influx in mBECs. We used a small glass capillary (exit diameter 12 μm) to apply a sub-cellular flow to mBECs, with a maximal surface shear of 2×10^4^ s^-1^, corresponding to a surface stress of 20 pN/µm^2^, approximately twice the mean value *in vivo* (Koller and Kaley, 1991). Bead tracers showed a nearly pencil-like laminar flow emerging from the pipette (Fig. S11). This flow clearly induced localized Ca^2+^ influx (Fig. 4D, *n* = 5 cells) and localized vesicle fusion (Fig. 4E, *n* = 4 cells) in the high-shear zones, without activating either mechanosensitive channels or vesicle fusion in other parts of the cell. This experiment establishes that localized changes in membrane tension drive sub-cellular signaling in a physiologically relevant context.

**Fig. 4.**
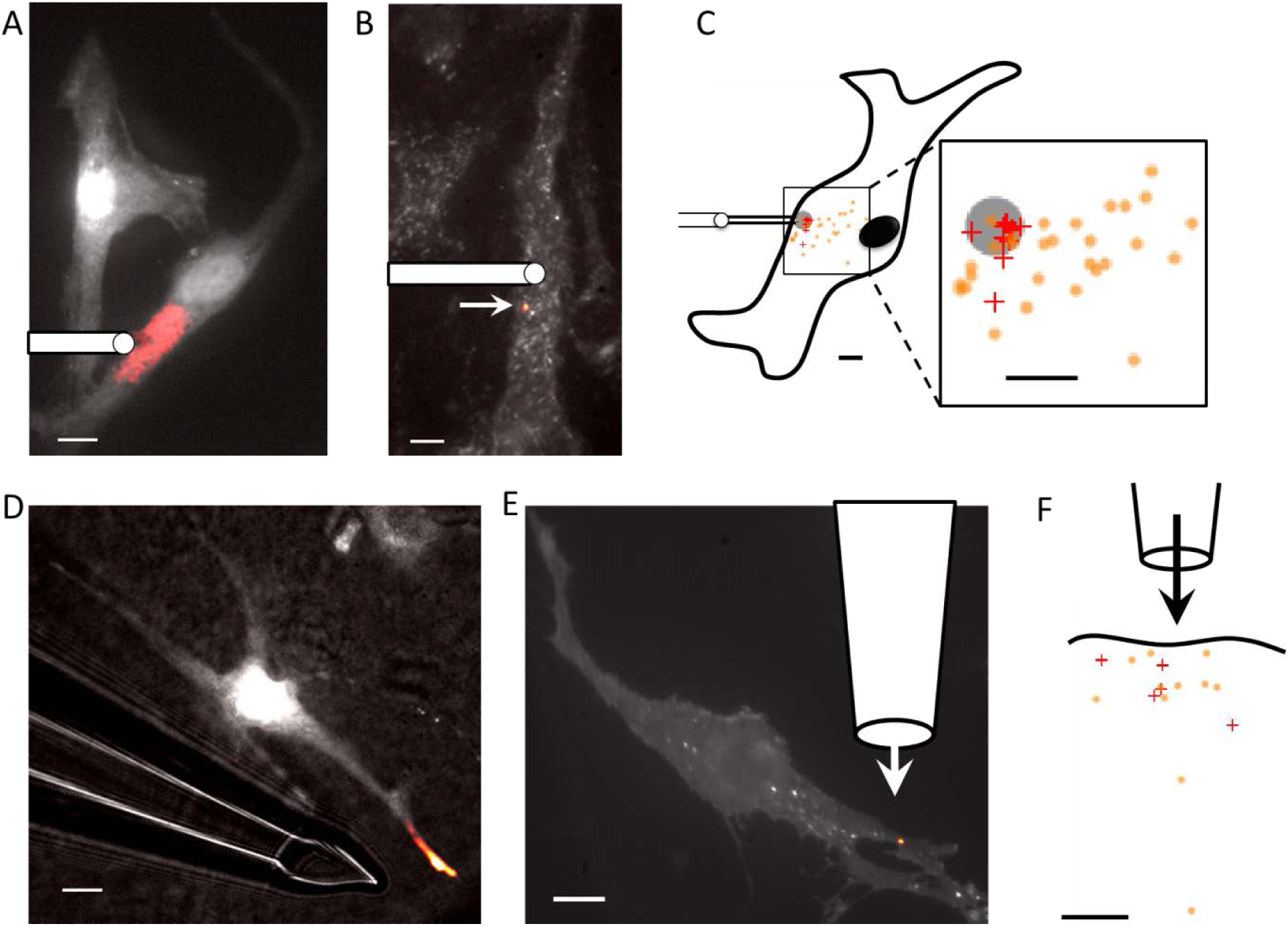
Tension mediates local activation of mechanosensitive ion channels and local vesicle fusion in primary mouse brain endothelial cells. **A)** Tether stretch triggered localized Ca^2+^ influx and **B)** vesicle fusion events. Images are composites of mean fluorescence (grey) and changes in fluorescence (heat map). Tether pulling pipette shown schematically. **C)** Distribution of Ca^2+^ influx (+) and vesicle fusion (o) sites relative to the tether attachment point (grey circle). Each mark represents one event (9 Ca^2+^ influx events from 7 cells; 29 vesicle fusion events from 6 cells). Average distance between Ca^2+^ initiation and tether attachment was 2.2 ± 0.5 μm (mean ± s.e.m), within the localization uncertainty (3 μm). Average distance between vesicle fusion and tether attachment was 8.0 ± 0.8 μm (mean ± s.e.m, vs. 28 ± 3 μm for null hypothesis). **D-E)** Local flow of extracellular buffer at 12 cm/s led to localized Ca^2+^ influx (**D**) and localized vesicle fusion (**E**). Images are composites of mean fluorescence (grey) and changes in fluorescence (heat map). In **D**, transmitted light shows the location of the pipette for flow delivery. **F)** Distribution of Ca^2+^ influx (+) and vesicle fusion (o) sites relative to the local flow. Each mark represents one flow-induced event (5 cells for Ca^2+^ influx; 11 fusion events from 4 cells for vesicle fusion). Scale bars in all panels 10 μm.

## DISCUSSION

Despite the well-established importance of membrane tension for many physiological processes (Basu et al., 2016; Boulant et al., 2011; Gauthier et al., 2011; Groves and Kuriyan, 2010; Houk et al., 2012; Keren et al., 2008; Maritzen and Haucke, 2017; Phillips et al., 2009; Ranade et al., 2015; Romer et al., 2007; Stewart et al., 2011), the path to equilibrium for local tension perturbations has received little attention (Raucher and Sheetz, 1999). Most studies have assumed that membrane tension is homogeneous across a cell (Diz-Muñoz et al., 2013; Gauthier et al., 2012; Gauthier et al., 2011; Houk et al., 2012; Keren et al., 2008; Keren, 2011; Morris and Homann, 2001; Mueller et al., 2017; Pontes et al., 2017; Watanabe et al., 2013). This assumption was justified either by analogy to isolated lipid bilayers (Keren et al., 2008; Watanabe et al., 2013), or by reference to experiments where membrane tension was globally perturbed via osmotic shocks or drug addition (Gauthier et al., 2011; Houk et al., 2012; Mueller et al., 2017; Raucher and Sheetz, 2000).

Our data and model provide direct evidence that there is no long-range propagation of membrane tension in cells, contrary to common belief. We showed that local perturbations to tension have local effects on signaling. Our findings also explain puzzling literature observations about the activation of PIEZO1 mechanosensitive ion channels in cells (Cox et al., 2016; Gottlieb et al., 2012; Lewis and Grandl, 2015): channel activation via pipette aspiration produced a current proportional to the area of the pipette aperture, not the whole cell area. This observation is easily understood if the increased tension is localized near the pipette.

Our study also suggests a resolution to a significant mystery in the field of mechanosensation. Direct measurements of resting cell membrane tension across various mammalian cells range from 3 to 40 pN/µm (Morris and Homann, 2001; Raucher and Sheetz, 1999), whereas the activation of mechanosensitive ion channels such as PIEZO and TREK channels requires a >100-fold higher membrane tension: 1000∼5000 pN/µm (Cox et al., 2016; Morris and Homann, 2001). Our study shows that a modest cell-wide stress can plausibly lead to a highly localized membrane strain, perhaps sufficient to activate MSCs. The cellular structures mediating such tension focusing remain to be identified.

A third important implication of our model is the extreme sensitivity of the tension diffusion coefficient *D*_σ_ to the area fraction of cytoskeleton bound obstacles, *Φ*_*i*_. The dramatic effect of ∼10% immobile obstacles on membrane rheology might seem counterintuitive. However, a similar effect is familiar in everyday life. An aqueous 10% collagen gel behaves as a solid and can be eaten with a fork. While the density of transmembrane obstacles has not been systematically studied, we anticipate that this important biophysical parameter will vary between cell types, between sub-cellular regions, and throughout the cell cycle. There may be physiologically important situations where tension can diffuse rapidly.

Changes in intracellular pressure could mediate long-range changes in membrane tension via the Laplace relation between pressure, membrane tension, and membrane curvature. Cytoplasmic pressure equilibrates quickly compared to membrane tension, but still more slowly than one might expect from bulk hydrodynamics (Charras et al., 2005). This effect was quantitatively explained by a poroelastic model in which intracellular proteins, membranes, and organelles formed an elastic gel through which pressure propagated diffusively, a three-dimensional analogue of the membrane model presented here. The diffusion coefficient of hydrostatic pressure was ∼10 µm^2^/s, more than 100-fold higher than that of membrane tension, reflecting the correspondingly lower viscosity of cytosol versus membrane (Kusumi et al., 2005).

Cytoskeletal reorganization could also mediate long-range changes in membrane tension (Bussonnier et al., 2014; Wu et al., 2013). One should think of the membrane and cytoskeleton as a composite material, in which deformation of the two components is tightly coupled. The far greater stiffness of the cytoskeleton compared to the membrane implies that the cytoskeleton dominates the rheology. In this composite picture, perturbations to the cytoskeleton could propagate quickly and cause long-range changes in membrane tension.

Many revisions to the fluid mosaic model have been proposed (Nicolson, 2014). Specialized structures, such as cytoskeletal corrals and lipid rafts, have been invoked to explain sub-cellular confinement in membranes (Kusumi et al., 2005). Indeed such structures are necessary to account for diffusional confinement and for local variations in membrane composition. Our results establish that a random array of transmembrane obstacles is sufficient to qualitatively change the membrane rheology from fluid to gel-like dynamics, without invoking any specialized structures. This mechanism of membrane gelation is unrelated to the lipid gel phases that arise at lower temperatures through phase transitions of the lipids themselves (Koynova and Caffrey, 1998). Within our model, the lipids remain liquid-like on the nanoscale, permitting free diffusion of molecular cargoes. Our model is entirely consistent with the thermodynamic data used to support the fluid-mosaic model (Nicolson, 2014; Singer and Nicolson, 1972), while adding a picture of the slow and heterogeneous approach to equilibrium.

## SUPPLEMENTAL INFORMATION

Supplemental Information includes Materials and Methods, Supplementary Discussions, 11 Supplementary Figures, and 1 Supplementary Table

## AUTHOR CONTRIBUTIONS

Z.S. and A.E.C. conceived the research, designed the experiments, analyzed the data and wrote the paper. Z.S. carried out the experiments. Z.T.G. carried out the experiment in Fig. S2. T.B. contributed to experimental design and data interpretation. H.A.S. guided the theoretical analysis.

## ACKNOWLEDGEMENTS

We thank Shahinoor Begum and Melinda Lee for help with neuron culture. We thank Katherine Williams, He Tian, Peng Zou, Yoav Adam, Linlin Fan, Sami Farhi, Veena Venkatachalam for help with molecular cloning and plasmid preparation. We thank Sean Buchanan from Lee Rubin’s Lab and Harry McNamara for providing primary mouse brain endothelial cells and giving advice on the culturing protocols. We thank Xiaowei Zhuang’s Lab for providing NIH 3T3 fibroblasts. We thank Guido Guidotti for helpful comments. This work was supported by the Gordon and Betty Moore Foundation and the Howard Hughes Medical Institute. Z.T.G. and T.B. were supported by NIH grant R01 GM 09755 and NIH grant U54CA193417. H.A.S. was supported by NSF grants CBET-1509347 and DMS 1614907.

## DECLARITION OF INTERESTS

The authors declare no competing financial interests.

## Supplementary Materials

### Materials and Methods

#### Cell culture, transfection, and staining

Henrietta Lacks (HeLa) cells, NIH/3T3 fibroblasts, and Madin–Darby canine kidney (MDCK) epithelial cells were cultured following standard protocols. Briefly, cells were grown in DMEM supplemented with 10% FBS and penicillin/streptomycin in a 37 °C incubator under 5% CO_2_. Cells were grown to 50-70% confluence in 3.5 cm dishes and transfected with 0.5 - 1 μg desired plasmid using TransIT-X2 (Mirus MIR6003). One day after transfection, cells were trypsinized and re-plated at a density of 10,000 - 30,000 cells/cm^2^ on glass-bottom dishes. Experiments were performed the following day. Before imaging, the cell culture medium was replaced with extracellular (XC) imaging buffer (125 mM NaCl, 2.5 mM KCl, 15 mM Hepes, 30 mM glucose, 1 mM MgCl_2_, 3 mM CaCl_2_, and pH 7.3). All lipids were purchased from Avanti Polar Lipids. Texas Red® DHPE was from Life Technologies.

All procedures involving animals were in accordance with the National Institutes of Health Guide for the care and use of laboratory animals and were approved by the Institutional Animal Care and Use Committee (IACUC) at Harvard. Hippocampal neurons from P0 rat pups were dissected and cultured in NBActiv4 medium at a density of 30,000 cells/cm^2^ on glass-bottom dishes pre-coated with poly-d-lysine (Sigma P7205) and matrigel (BD biosciences 356234). At 1 day in vitro (DIV), glia cells were plated on top of the neurons at a density of 7000 cells/cm^2^. At DIV5 - 7, neurons were transfected following the calcium phosphate protocol (Jiang and Chen, 2006). Imaging was performed 5 - 7 days after transfection, with neuron culture medium replaced with XC buffer.

Primary mouse brain endothelial (mBEC) cells were dissected and cultured in complete mouse endothelial cell medium (Cell Biologics M1168). For tether imaging or Ca^2+^ imaging, cells were plated at a density of 10,000 - 30,000 cells/cm^2^ on glass-bottom dishes and stained with CellMask™ (Thermo Fisher C37608) for 10 minutes or with Fluo-4-AM (Life Technologies F14201) for 30 min before experiments. For vesicle imaging, cells were grown to 50% confluence in 3.5 cm dishes and transfected with lenti-virus encoding mOrange2-TM. 5 - 7 days after transfection, cells were trypsinized and re-plated at a density of 10,000 - 30,000 cells/cm^2^ on glass-bottom dishes. Experiments were performed 12 – 36 hours after cells were plated to glass-bottom dishes. Before imaging, the cell culture medium was replaced with XC buffer. Neurons and mBEC cells were fed twice weekly until experiments.

For nonspecific extracellular staining of transmembrane proteins, cells were incubated with 250 μg/mL Alexa Fluor™ 488 NHS Ester (Thermo Fisher A20000, dissolved using the original cell culture medium) for 30 minutes. Cells were then washed 3 - 5 times with 1 mL XC buffer before imaging. Amaranth (Sigma 87612), with a final concentration of 500 µM, was used to quench the Alexa488 fluorescence.

For imaging intracellular vesicles with FM 4-64 (Thermo-Fisher T13320), cells were incubated with 20 μg/mL FM 4-64 for 20 minutes. Cells were then washed 5 times with 1 mL XC buffer before imaging, leaving the cell with only intracellular vesicles stained. Fusion of vesicles was reported as the disappearance of fluorescent puncta (Gauthier et al., 2009). Ionomycin (Sigma I9657), with a final concentration of 5 µM, was used to trigger cell-wide vesicle fusion.

#### DNA constructs

pCAG: GPI-eGFP (Addgene plasmid # 32601, eGFP targeted to the plasma membrane using the glycophosphatidylinositol anchor) was a gift from Anna-Katerina Hadjantonakis (Rhee et al., 2006). DRD2-eGFP: dopamine receptor D2 with eGFP. CheRiff-eGFP: an ultra-sensitive, fast, and well trafficked channelrhodopsin variant linked with eGFP (Hochbaum et al., 2014). ASAP1: Accelerated Sensor of Action Potentials 1 (St-Pierre et al., 2014). eGFP-TM: eGFP targeted to the extracellular side of the plasma membrane, using a transmembrane helix from PDGF receptor on the pDisplay™ Mammalian Expression Vector (Thermo Fisher). eGFP-KRAS: eGFP targeted to the inner leaflet of plasma membrane using the C-terminus sequence of KRAS. R-CaMP2 was a gift from Haruhiko Bito (Inoue et al., 2015). GCaMP6f (Addgene plasmid # 40755) was a gift from Douglas Kim (Chen et al., 2013). PIEZO1-mCherry was a gift from Ardem Patapoutian. mOrange2-TM: mOrange2 targeted to the extracellular side of the plasma membrane, using a transmembrane helix from PDGF receptor on the pDisplay™ Mammalian Expression Vector.

#### Bleb formation

Blebs were induced by treating the cells grown on glass bottom dish with 100 - 200 μM latrunculin B (Sigma L5288) dissolved in 200 μL XC buffer. Blebs started forming within 3 minutes of drug addition. Then, 2 mL of XC buffer was added to the dish and majority of the blebs became stable for further experiments.

#### Glass micropipette fabrication, tether pulling, and imaging

Micropipettes were pulled from glass capillaries (WPI 1B150F-4) using a pipette puller (Sutter Instrument P1000). The tip of the pipette was cut to an opening diameter of ∼3 μm and bent to ∼40° using a microforge (WPI DMF1000). Experiments were performed on a home-built epi-fluorescence microscope (Kralj et al., 2011). Two Sutter manipulators (Sutter Instrument MP-285) controlled pipette motion.

The pipettes were immersed in a dispersion of 4 μm diameter Anti-Digoxigenin coated polystyrene beads (Spherotech DIGP-40-2), and suction was applied to plug each pipette aperture with a single bead. The beads were then brought into contact with cell membranes and retracted to pull out membrane tethers. A digital micromirror device (DMD) with 608 × 684 pixels (Texas Instruments LightCrafter) patterned the illumination to confine fluorescence excitation to the tether regions. In cases where tethers broke, the piece of tether attached to cells retracted to its mother cell within one minute. To obtain large membrane tension changes on blebs through tether pulling, it is advantageous to choose more spherical blebs. Otherwise, on floppy blebs, a change of tether length for ∼ 100 μm does not result in measurable changes in tether fluorescence (or equivalently, membrane tension).

Measurements of tension-dependent tether pulling force on GUVs (Fig. S1) were performed on a home-built optical trap with 6 μm diameter streptavidin coated beads (Polysciences, Inc) as described previously (Heinrich et al., 2010; Shi and Baumgart, 2015). Calibration between tether pulling force and tether intensity (with HeLa cells, as shown in Fig. S2C and S2D) was achieved with simultaneous recording of pulling force (through the optical trap) and tether fluorescence (with patterned illumination).

Tether diameters were estimated by imaging HeLa cells expressing membrane-bound fluorescent proteins. By measuring the cumulative fluorescence *I*_cell_ in a patch of flat cell membrane of area *A*_cell_ and the cumulative fluorescence *I*_tether_ on tethers (pulled from the same cell) of length *l* (Fig. S2B), the diameter of a single tether is calculated using:

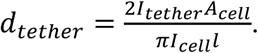

The factor of 2 in the numerator accounts for the fact that the cell has top and bottom membranes, both of which contribute to *I*_*cell*_. All experiments with cells were performed with a 60x oil objective (Olympus UIS 2, N.A. 1.49) with an objective heater (Bioptechs) to keep the sample at 37 °C.

#### Fluorescence recovery after photobleaching (FRAP) measurements of diffusion

To measure tracer diffusion on cell membranes, a flat patch of membrane was photobleached within a circular region of radius *r* = 7 μm at an illumination intensity of 1 kW/cm^2^ for ∼ 60 s. Then an illumination intensity of 0.5 W/cm^2^ was used to monitor the recovery. To measure tracer diffusion on tethers, the same laser spot was used to bleach a *d* = 14 μm long region of tether. Measurements on plasma membrane and tether were performed sequentially on the same cells.

For FRAP measurements on the cell, the fluorescence recover was fit to the relation:

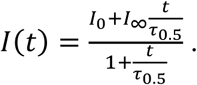

The diffusion coefficient was extracted using (Kang et al., 2012) 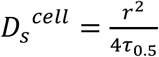.

For FRAP measurements on the tether, the recovery phase was fit to:

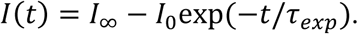

The diffusion coefficient was extracted using (Rosholm et al., 2017) 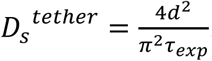. The high membrane curvature in tethers is reported to slightly decrease diffusion relative to a planar bilayer (by less than a factor of 2) (Domanov et al., 2011), an effect that we neglected.

#### FRAP measurement of fraction of transmembrane proteins that are immobile

To measure the immobile fraction of cell surface proteins, NHS-ester Alexa 488 labeled cells were bleached with a donut shape laser beam (inner radius 17.5 μm, outer radius 35 μm) following the procedure described above.

The immobile fraction of proteins was calculated using

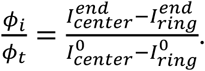

Here, the *ϕ*_*t*_ represents the area fraction of all labeled transmembrane proteins. 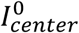 and 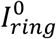 represent fluorescence intensities right after photobleaching in unbleached and bleached regions respectively (at *t* ∼ 100 s, see Fig. S6C). 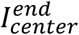 and 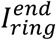 represent fluorescence intensities, in unbleached and bleached regions respectively, at the end of the FRAP experiment (t = 1000 s).

#### Simulation of relaxation of membrane tension in a tether

To model tension relaxation in a pulled tether, we decomposed the experiment into three steps:

##### (I) Initial equilibrium

Tether length *l*, radius *r* and membrane tension, *σ*, are related by:

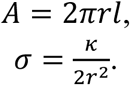

Combining these two relations leads to an expression for membrane tension as a function of tether length and area:

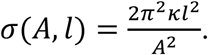

##### (II) Elongation of tether at constant velocity, v_pull_

The simulation is broken into small time-steps of length *Δt* = 3 s. For each time-step, the dynamics are described by three processes:

1. At constant tether area, *A*, increase tether length by *Δl* = *ν*_*pull*_*Δt*. Then update *l, r*, and *σ*, without allowing flow of lipids from the cell into the tether:

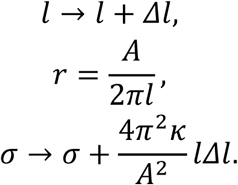
2. Calculate the diffusion of tension in the cell membrane, treating the membrane as a 20 µm radius disk and matching tension across the cell-tether boundary at radius *r*:

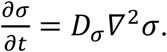
3. Calculate flow of lipids from the cell membrane into the tether, keeping *l* constant. The flux into the tether is given by the solution to the tension diffusion equation at the cell-tether boundary:

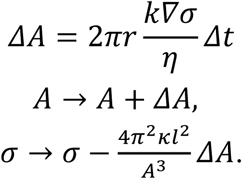

##### (III) Relaxation of tension via lipid flow into a tether of constant length

The steps are the same as in (II) except that tether length is always kept constant. The results of these simulations are plotted in Fig. S7.

#### Tether pulling and Ca^2+^ imaging

For simultaneous imaging of tethers and Ca^2+^ influx, MDCK cells were co-transfected with GPI-eGFP and R-CaMP2. Blue laser light (488 nm) for exciting GPI-eGFP was confined to the tether region via a digital micromirror device, while green laser light (532 nm) for exciting R-CaMP2 illuminated the whole cell (Fig. 3B). Images were acquired continuously at 5 Hz with an emission filter simultaneously passing GFP and RFP emission wavelengths. Initiation points of Ca^2+^ influx were determined as the center of the Ca^2+^ influx (as shown in the heat map of Fig. 3C) in its first observable frame.

In the studies of the effects of Gd^3+^ and GsMTx4 (Fig. 3D) on Ca^2+^ influx, MDCK cells were transfected with GCaMP6f as a Ca^2+^ reporter. Under wide-field 488 nm excitation, images were acquired continuously at 2 Hz with an emission filter for GFP.

In the study of the effect of PIEZO1 on Ca^2+^ influx, MDCK cells were co-transfected with PIEZO1-mCherry and GCaMP6f. PIEZO1-mCherry expressing cells were identified with 532 nm excitation and an emission filter for RFP. Images were acquired continuously at 2 Hz under wide-field 488 nm excitation with an emission filter for GFP.

All tether pulling experiments shown in Fig. 3D and 3I followed the same tether pulling protocol (move the bead to gently touch a GCaMP6f expressing cell for 20 s, then pull bead away from the cell for 500 µm with the first 200 µm at 5 µm/s and the next 300 µm at 10 µm/s). Changes of GCaMP6f fluorescence (F_max_/F_0_) were measured in the region of bead-cell attachment (diameter 4 µm circle) with corrections for background and photo bleaching. F_max_ is the peak fluorescence during tether elongation; F_0_ is the fluorescence baseline before tether pulling. We poked holes on the cells at the end of each tether pulling experiment to verify that the cells could report Ca^2+^ influx. In mBEC cells, the same tether pulling protocol was applied to cells stained with Fluo-4-AM (Fig. 4A).

#### Tether pulling and vesicle fusion

The same tether pulling protocol used for MSC activation and Ca^2+^ imaging was applied to cells expressing mOrange2-TM (Fig. 3G and 4B). Under wide-field 532 nm excitation, images were acquired continuously at 1 Hz with an emission filter for YFP. Vesicle fusion sites were determined as the center of bright dots that appeared during pulling (as shown in the heat map of Fig. 3G and 4B).

To determine the distribution of tether-vesicle distances expected from the null hypothesis (vesicles fuse at random locations in the cell), the image of each cell was converted into a binary mask. The mean distance from the tether location to all points on the cell was then calculated. This distance was averaged over all cells measured.

#### Local flow experiments with mBEC cells

mBEC cells were stained with Fluo-4-AM or transfected with mOrange2-TM as described above. Pipettes with an exit diameter of *R*_p_ = 12 µm were used to inject XC buffer near one end the cell by quickly increasing the pressure inside pipette. Fluorescence images were acquired at 2 Hz. The flow speed was calibrated by measuring the rate of decrease of buffer volume in the pipette. For data shown in the main text, this rate was *Γ* = 14 nL/s. Buffer speed exiting pipette was calculated by *v*_flow_ = *Γ*/ (π*R*_p_^2^) = 12 cm/s. Maximal surface shear induced by the pipette was approximately *v*_flow_/*R*_p_ = 2×10^4^ s^-1^, corresponding to a surface stress of η_c_·*v*_flow_/*R*_p_ = 20 pN/µm^2^. Here, η_c_ = 10^-3^ Pa·s is the viscosity of XC buffer.

### Supplementary Discussion

#### Relation between tether radius, pulling force, and membrane tension

For a tube of length *l* and radius *r*, the free energy of a tether *U* is (Brochard-Wyart et al., 2006; Derényi et al., 2002):

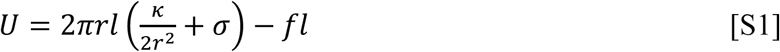

Here *κ* is the bending stiffness of the membrane, *σ* is membrane tension, and *f* is the external pulling force.

The surface tension acts to reduce the radius (and therefore decrease the total area of the tether) while the bending stiffness works to increase the radius (to avoid membrane bending). The balance between these two forces sets the mechanical equilibrium. The equilibrium radius *r*_0_ and pulling force *f*_0_ are obtained from:

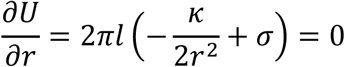

and

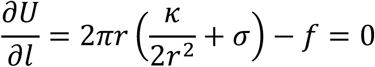

from which we obtain:

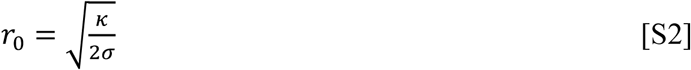

and

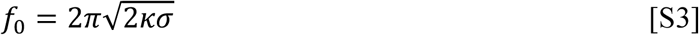

These equations show that one can determine the tension of a bilayer by measuring either the tether radius or the pulling force.

#### Hydrodynamics of lipid flow

To describe lipid flow through a medium comprised of randomly dispersed immobile obstacles, the Stokes equation is augmented with a drag term (Bussell et al., 1995; Howells, 1974) 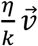:

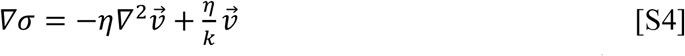

where *σ* is the membrane tension, *η* is the two-dimensional membrane viscosity, 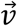 is the velocity field of lipid flow, and *k* is the Darcy permeability of the array of obstacles. When the Brinkman equation is written for pressure, rather than membrane tension, the signs on the right-hand side are reversed (fluids flow from high to low pressure, but membranes flow from low to high tension). Physically, the ratio 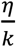 is the drag coefficient of a fixed array of obstacles.

Conservation of mass requires that:

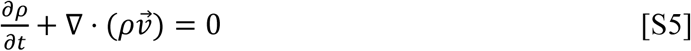

where *ρ* is the two-dimensional density of lipids. Assuming a small perturbation to lipid density, *ρ* = *ρ*_*o*_ + *δρ*, Eq. S5 becomes:

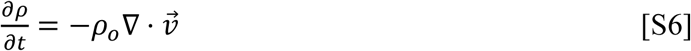

We assume a linear stress-strain relation for the membrane tension (Hochmuth et al., 1973):

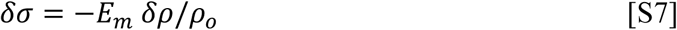

where *E*_*m*_ is the effective area expansion modulus of the cell membrane. Equations S4, S6, and S7 describe the hydrodynamics of lipid flow in cell membranes containing random arrays of fixed obstacles. The equations can be combined to obtain:

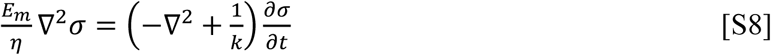

Neglect of membrane obstacles is equivalent to keeping only the first term on the right-hand side of Eq. S4 or S8.

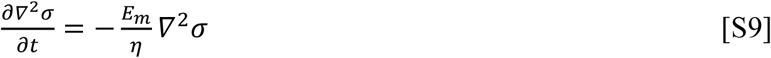

This relation identifies a relaxation time for tension fluctuations, *τ* = *η*/*E*_*m*_. Inserting estimates of membrane viscosity (*η* = (3.0 ± 0.4) × 10^-3^ pN·s/μm) and area expansion modulus (*E*_m_ = 40 pN/μm)(Hochmuth, 2000) gives a relaxation time less than 0.1 ms, as has been used in the literature (Keren et al., 2008).

If the spacing between transmembrane obstacles is small compared to externally imposed variations in the flow field, then the second term of Eq. S4 or S8 dominates, and we obtain a diffusion-like equation for membrane tension:

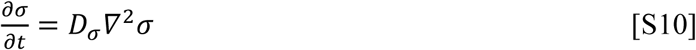

with 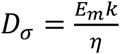 In other words, if *l*_*c*_ is a characteristic length scale of variations in *σ* then Eq. S10 applies when 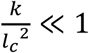. An essentially identical result describes the three-dimensional propagation of pressure in a porous elastic medium (Charras et al., 2005).

#### Calculation of drag due to a random array of fixed cylinders

When *N*_*i*_ immobile proteins are present in a piece of membrane with area *A*, the immobile area fraction is 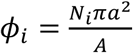, where *a* is the radius of one immobile particle. Bussel *et al.*(Bussell et al., 1995) adapted Howell’s mean-field solution of the Stokes equation for a random array of cylindrical obstacles (Howells, 1974) to calculate the mean force, 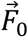, required to drag one particle with a mean velocity 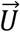 through the background of immobile particles.

We introduce the dimensionless quantity 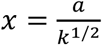, where *a* is the radius of an obstacle and *k* is the Darcy permeability coefficient. Then:

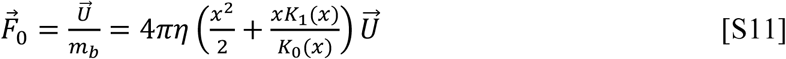

Here, *m*_*b*_ is the mobility of the tracer particle, and *K*_*0*_ and *K*_*1*_ are the modified Bessel functions of the second kind with orders of 0 and 1, respectively.

According to the Brinkmann model, the mean force per unit area to drag the membrane at velocity 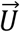 is:

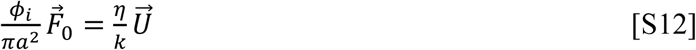

Together, these two equations lead to the relation between permeability and immobile protein fraction:

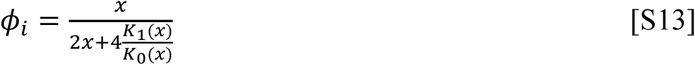

The dimensionless diffusion coefficient of tracer molecules is related to the mobility via:

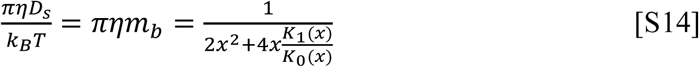

The dimensionless diffusion coefficient of membrane tension is:

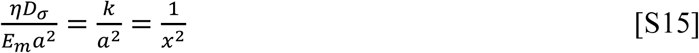

The upper limit of tracer diffusion is given by the Saffman–Delbrück model (Saffman and Delbrück, 1975):

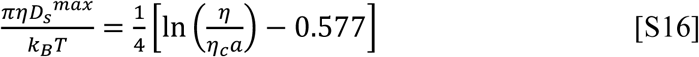

where *η*_*c*_ is the viscosity of the fluid medium surrounding the membrane and it is assumed that 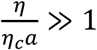.

For diffusion of tension, hydrodynamic coupling to the cytoskeleton exerts viscous drag even in the absence of fixed obstacles. This drag sets an upper bound on the diffusion coefficient for membrane tension:

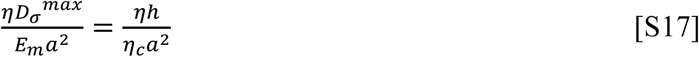

where *h* is the distance between the cortical cytoskeleton and the plasma membrane (see Fig. 2). We used *η*_*c*_= 10^-3^ pN·s/μm^2^, *a* = 2 nm (Bussell et al., 1995), and *h* = 20 nm (Clausen et al., 2017).

#### Equation of state of membranes

The relation between the projected membrane density and the membrane tension is given in equation [S7]. The resting tension of the cell membrane (∼25 pN/µm) is much lower than the tension where enthalpic stretching of lipid bilayers becomes significant (500 pN/µm) (Evans and Rawicz, 1990). Microscopically, the membrane has undulations due to thermal fluctuations, due to fluctuations in the underlying cytoskeletal support, and possibly due to binding of curvature-inducing proteins (Shen et al.) (e.g. in caveolae). The membrane density, *ρ*_0_, refers to a projected density of lipids after averaging over these microscopic undulations. Tension can partially smooth these undulations, leading to an effective stretching modulus that does not involve changing the mean spacing between lipid molecules. Experimental measurements of the effective area expansion modulus of cells (Hochmuth, 2000) indicate that the apparent elasticity is mainly associated with structures such caveolae and microvilli and the contribution from thermal agitation is negligible (Evans and Rawicz, 1990; Hochmuth et al., 1973; Hochmuth, 2000).

## Supplementary Figures

**Fig. S1.**
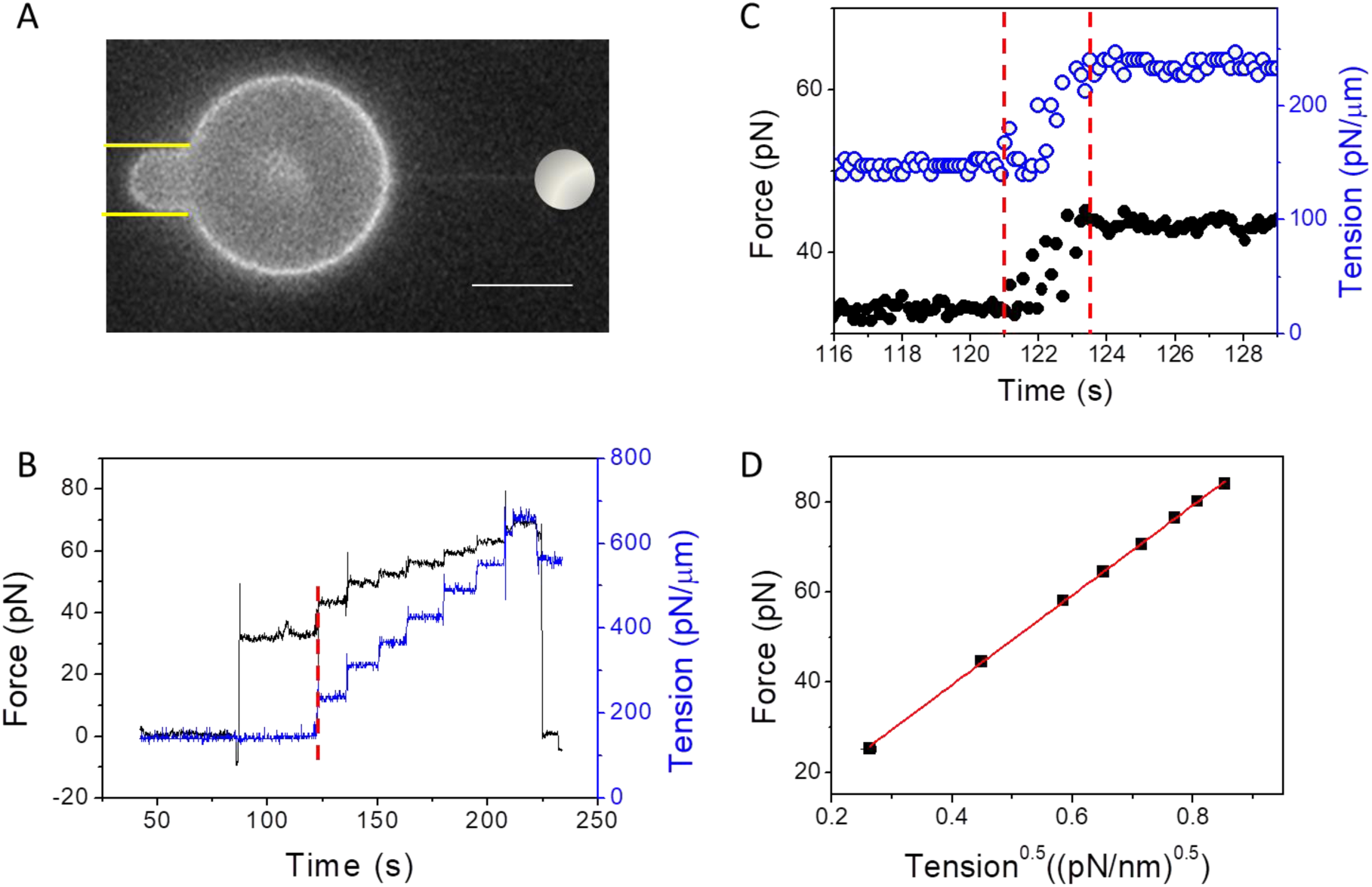
Membrane tension equilibrates quickly in artificial lipid bilayers. **A)** Fluorescence image of a micropipette-aspirated GUV (DOPS:DOPE:DOPC = 35:30:35) containing 0.5% DSPE-Bio-PEG2000 and labeled with 0.3% Texas Red® DHPE. The edge of the pipette is marked with yellow lines. An optically trapped bead (white circle) pulled a membrane tether opposite to the pipette. Scale bar 10 μm. **B)** Changes in membrane tension (blue) were induced by applying steps of pressure to the pipette and tether pulling force (black) was monitored via the optical trap. **C)** Close-up of the step marked with a red line in **B**, showing no detectable delay between change in tension and change in tether force. Measurements sampled at 10 Hz. **D)** Relation between tether pulling force and the square root of membrane tension, with a linear fit following the relation: 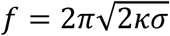 (red line, R^2^ = 0.99). Error bars are s.e.m..

**Fig. S2.**
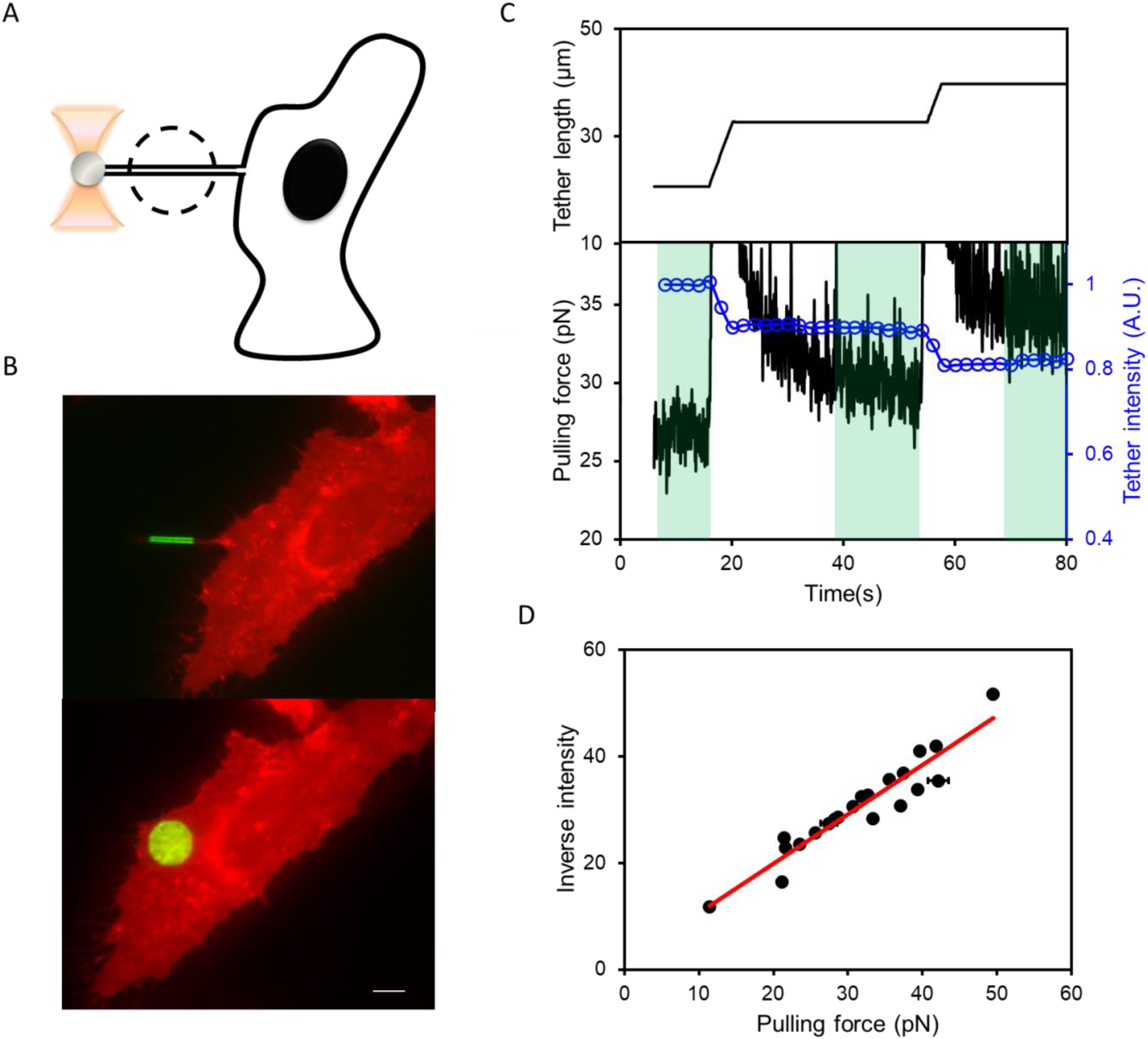
Tether diameter reports membrane tension. **A)** Schematic showing simultaneous measurement of tether pulling force (with an optical trap) and tether fluorescence intensity (with patterned illumination in the dashed circle). **B)** Determination of tether diameter from tether fluorescence intensity. A 14 µm diameter circular spot of illumination was first directed to a tether (top) and then to a flat portion of the parent cell (bottom). The ratio of the total fluorescence excited in these two configurations equals the ratio of illuminated membrane areas. These calibrations yielded an average tether diameter *d*_tether_ = 150 ± 10 nm (mean ± s.e.m., n = 5 cells) for tethers ∼20 μm long. Scale bar, 10 μm. **C)** Tethers were pulled with a bead in an optical trap, and tether length, fluorescence intensity, and force were measured simultaneously. Regions shaded in green were used to calculate the relation between steady state tether fluorescence intensity and pulling force. **D)** Relation between tether pulling force and inverse of tether fluorescence intensity (normalized to expression level; R^2^ = 0.9, n = 7 cells). Error bars are s.e.m. Experiments were performed on HeLa cells at 37 °C expressing membrane-targeted fluorescent protein mOrange2-KRAS.

**Fig. S3.**
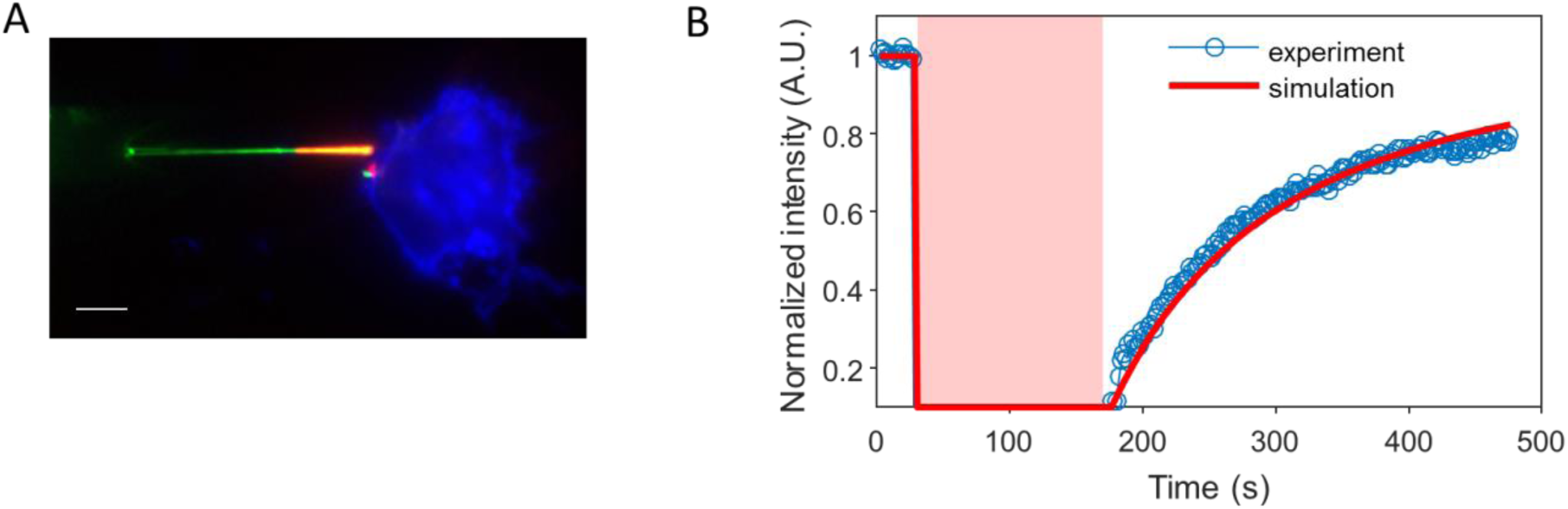
Tethers are in diffusive contact with the plasma membrane. **A)** Composite fluorescence image showing the photobleached region (red) on a tether (green), attached to a HeLa cell (blue) expressing DRD2-eGFP. Scale bar, 10 μm. **B)** FRAP of the tether (blue) and corresponding simulation (red) assuming free diffusion from cell to tether. The simulation used the experimentally measured diffusion coefficient of DRD2-eGFP on the cell, D_s_(Cell) = 0.037 µm^2^/s (see Table S1), and a tether radius of 75 nm. The simulation was performed in Matlab. Time of photobleaching is shaded red.

**Fig. S4.**
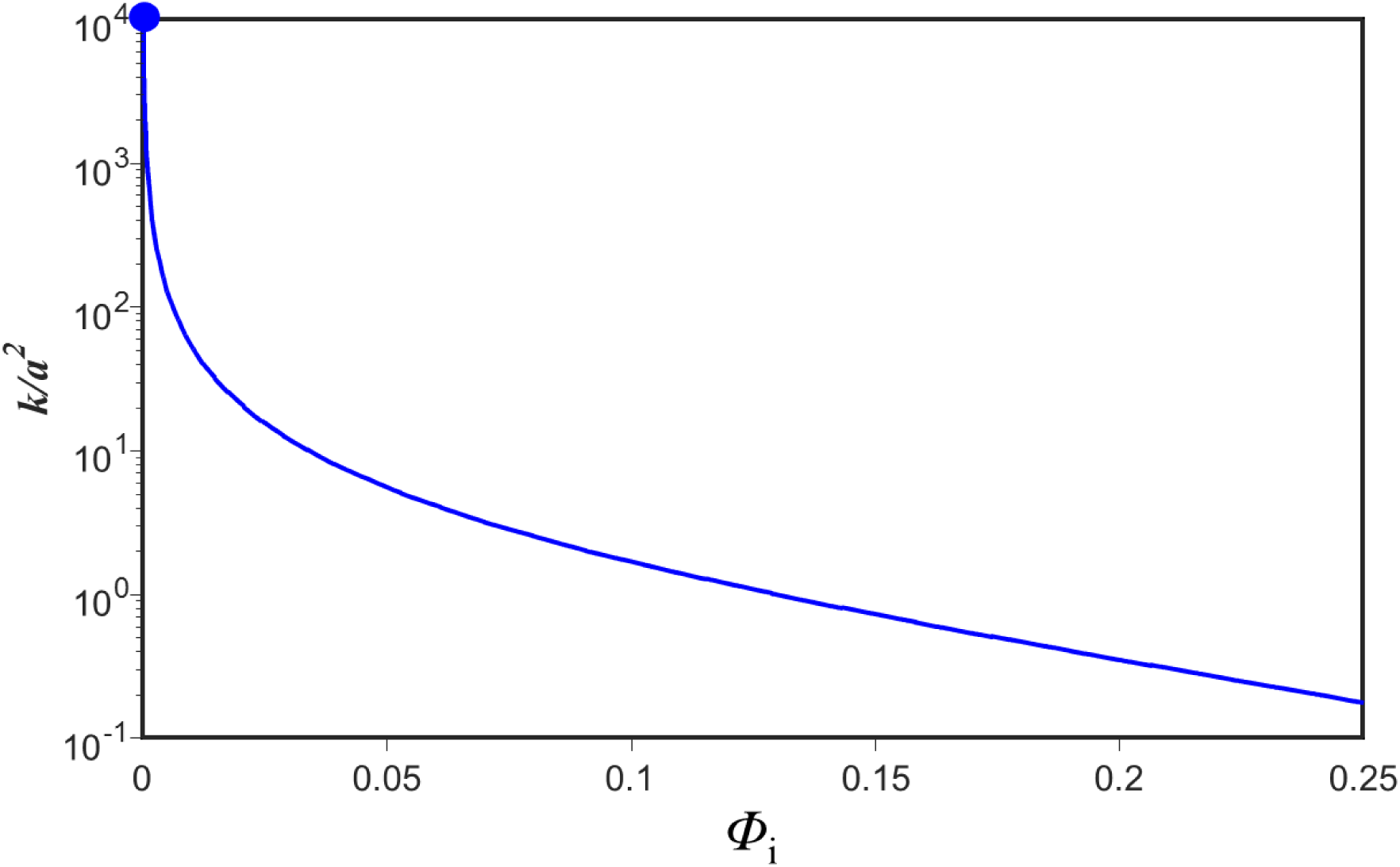
Relation between Darcy permeability *k* and area fraction of immobile proteins *Φ*_*i*_. The function 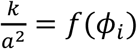 was derived by Bussell (Bussell et al., 1995) *et al.*, who showed that 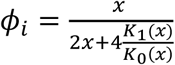, where 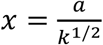 see *Supplementary Discussion*). The upper limit (blue dot) is calculated from viscous drag of the cytoplasm layer between membrane and actomyosin cortex

**Fig. S5.**
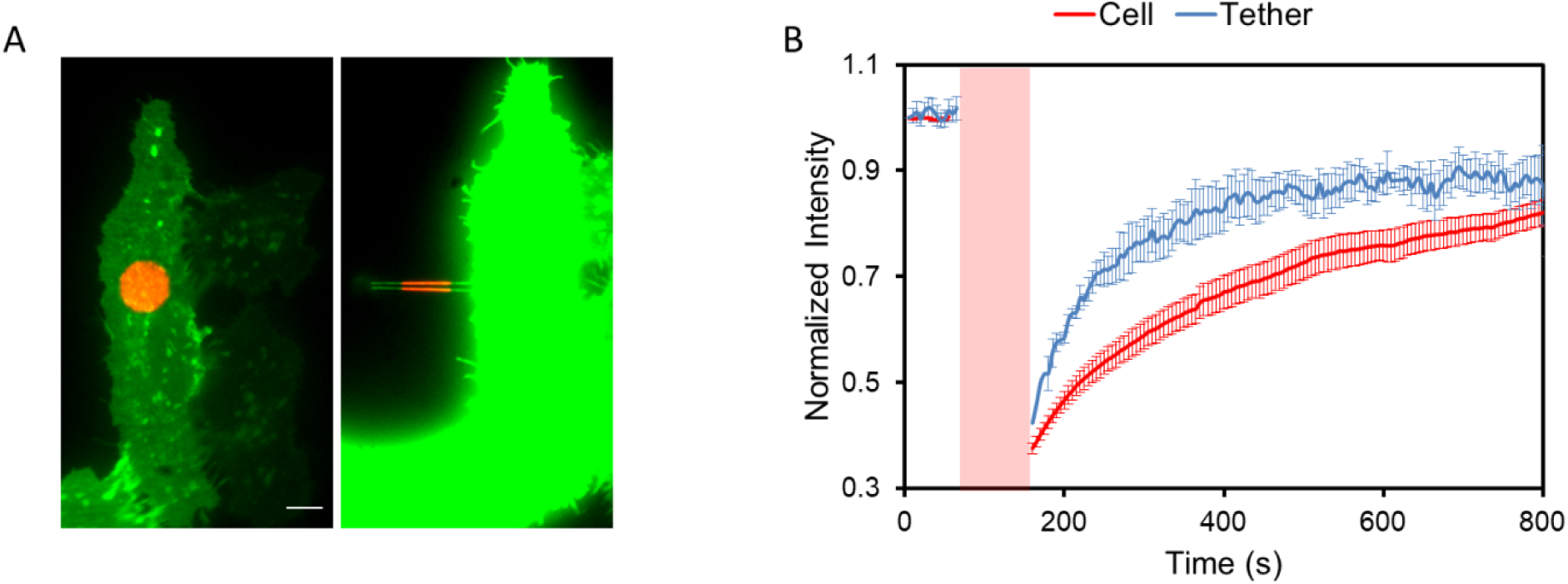
FRAP measurements of tracer diffusion on cell membranes and on tethers. **A)** Composite images showing the cell (green) and 14 µm circular photobleaching spot (red). Left: FRAP on cell membrane. Right: FRAP on tether. Scale bar, 10 μm. **B)** FRAP data in HeLa cells expressing DRD2-eGFP. Error bars represent s.e.m. from *n* = 10 tethers pulled from 10 cells. Time of photobleaching is shaded red.

**Fig. S6.**
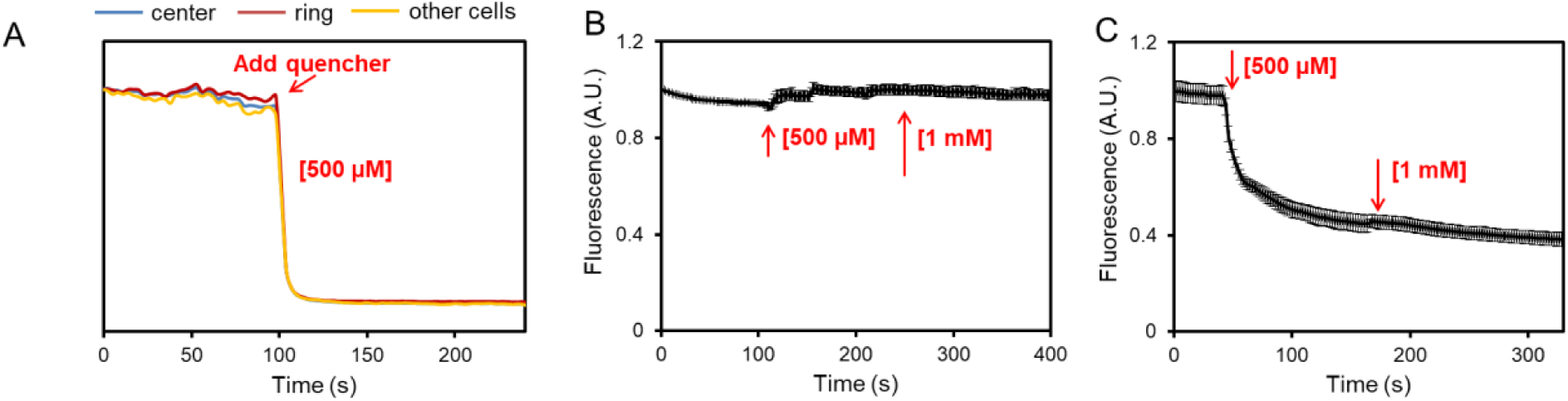
Validation of FRAP measurement of the immobile fraction of transmembrane proteins. **A)** After the FRAP measurement, a cell impermeant fluorescent quencher, amaranth, was added to a final concentration of 500 µM to quench the Alexa488 fluorescence. Fluorescence of all regions of the cell membrane dropped to background levels, establishing that there was no detectable internalization of fluorescently labeled proteins. **B-C)** Validation that amaranth functions as a cell-impermeant fluorescence quencher. **B)** In cells expressing, intracellular eGFP (eGFP-KRAS) amaranth did not affect fluorescence but **C)** in cells expressing extracellular eGFP (pDisplay: eGFP-TM) amaranth quenched fluorescence. Error bars are s.e.m. over *n* = 4 cells for eGFP-KRAS and *n* = 5 cells for eGFP-TM.

**Fig. S7.**
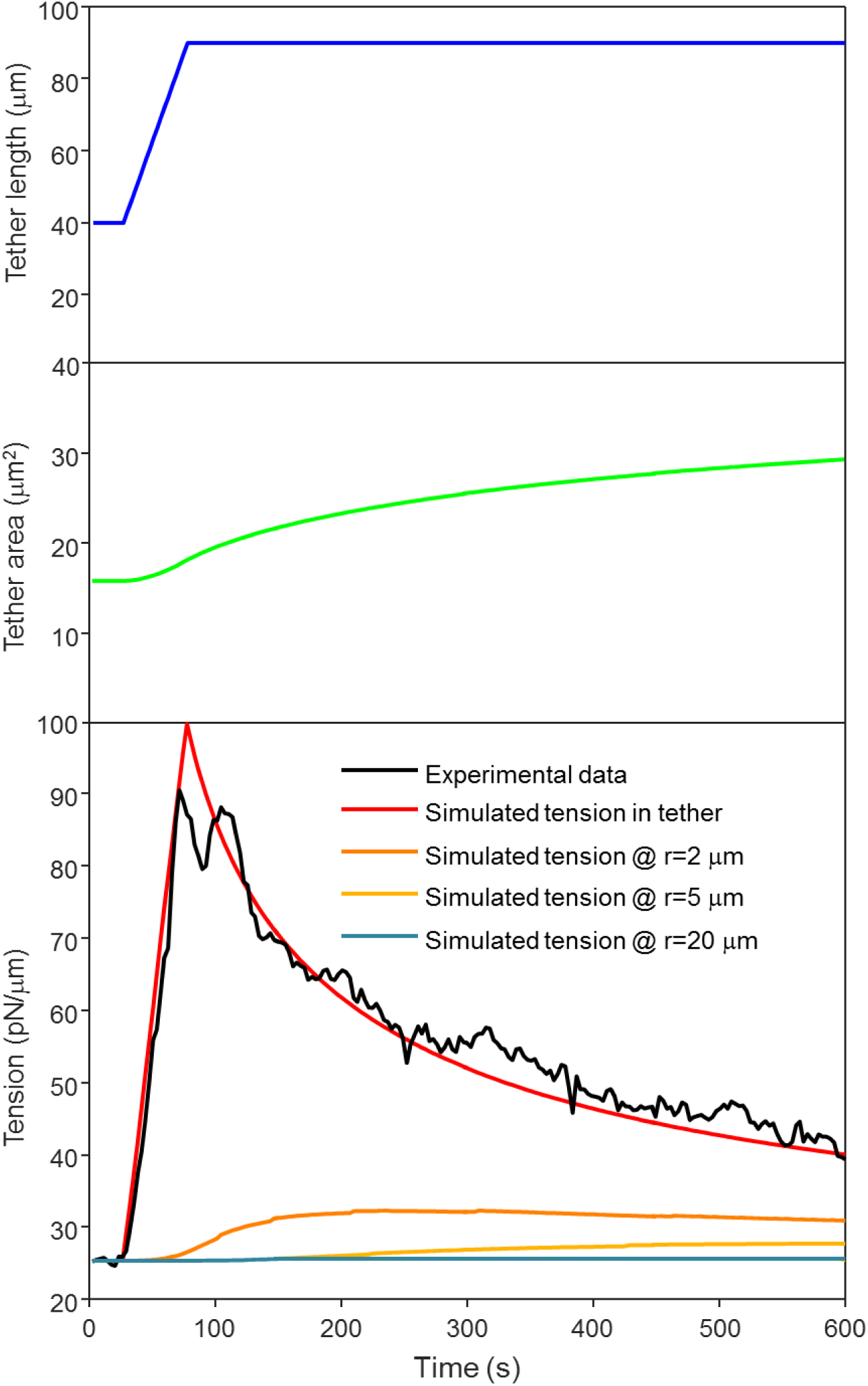
Simulation of time-dependent pulling force in a membrane tether. Parameters were: effective area expansion modulus in the cell-attached plasma membrane *E*_m_ = 40 pN/μm; tension diffusion coefficient *D*_σ_ = 0.024 μm^2^/s. The simulation matched the tether pulling experiment shown in **Fig. 1G** (initial tension σ_0_ = 25 pN/μm, ramp increase in tether length from 40 μm to 90 μm at a pulling speed *v*_pull_ = 1 μm/s). All simulations were done with Matlab. See *Materials and Methods* for details of the simulation.

**Fig. S8.**
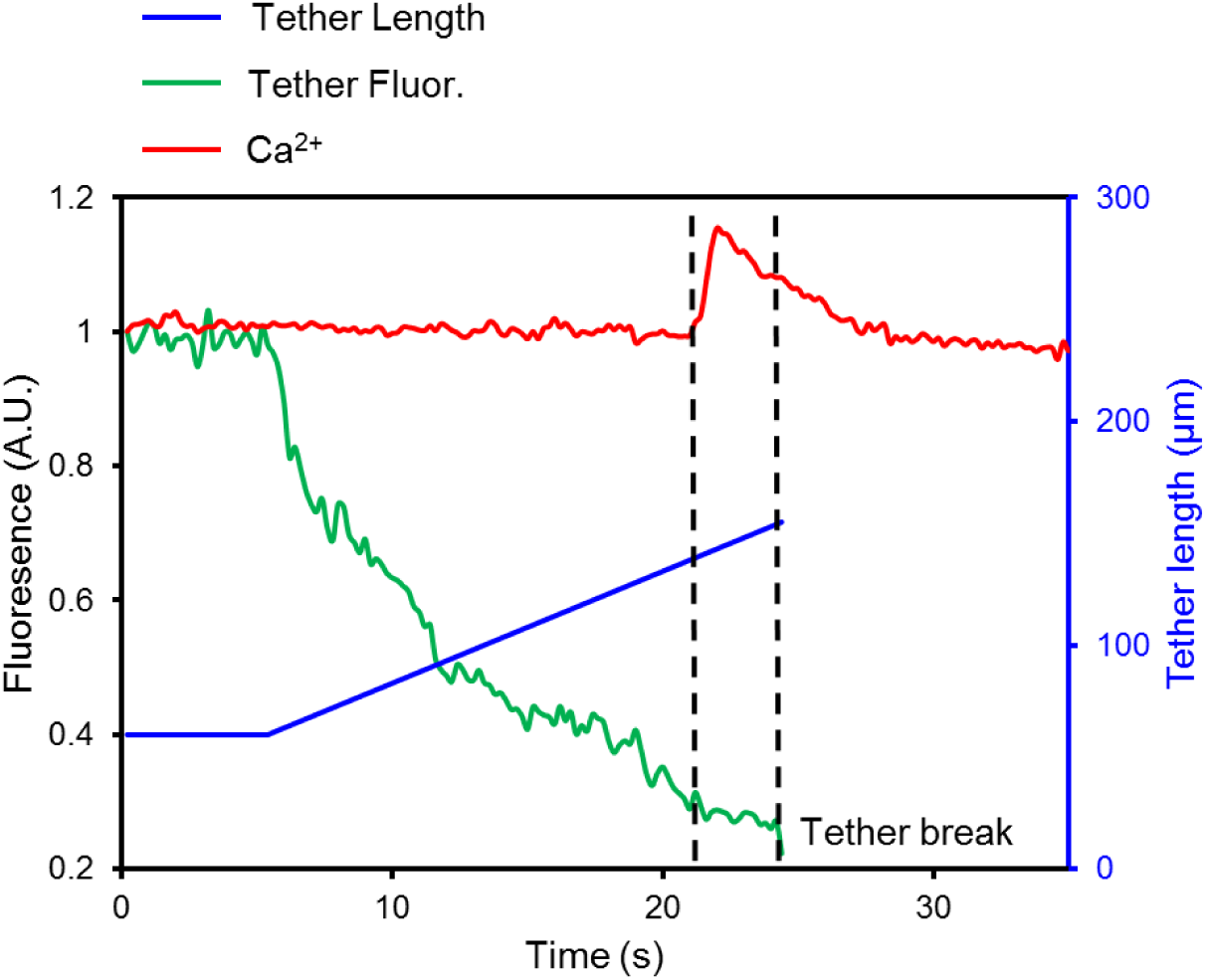
Simultaneous imaging of tether fluorescence and intracellular Ca^2+^ during tether pulling experiments. In experiments as shown in Fig. 3b, tether fluorescence intensity reports tether radius. Membrane tension can then be estimated from 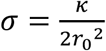 (see *Supplementary Discussions*). The activation tension of MSCs was found to be approximately 11 times higher than the resting tension of the cell (dashed line, resting membrane tension is ∼25 pN/µm). This estimate of activation tension is a lower bound because at the time of MSC activation the tether was still elongating, so tether diameter was not fully equilibrated.

**Fig. S9.**
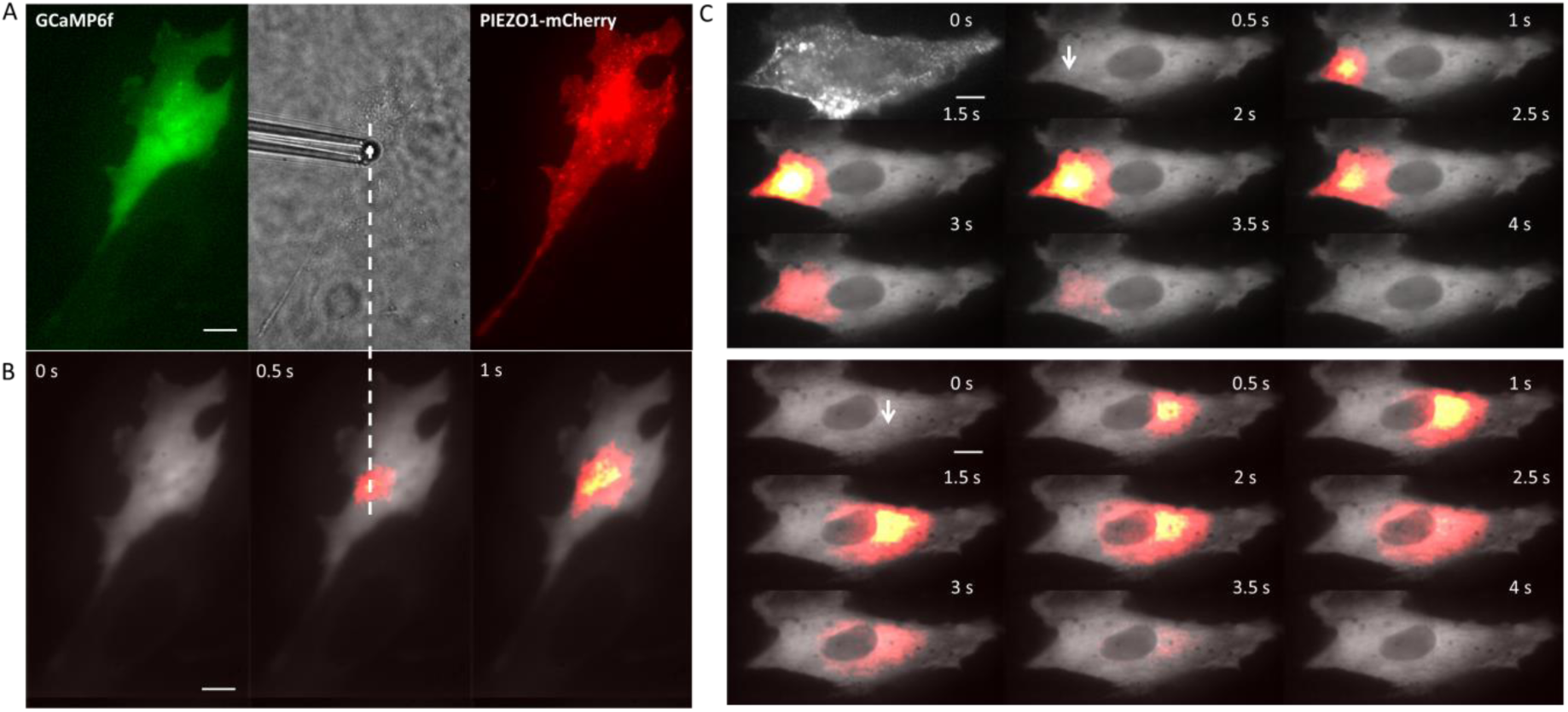
Local activation of mechanosensitive ion channels in MDCK cells expressing PIEZO1-mCherry. **A)** MDCK cell co-expressing GCaMP6f (green) and PIEZO1-mCherry (red), with a pipette-controlled bead locally touching the cell (grey). **B)** Localized Ca^2+^ influx triggered by tether stretch. Dashed line shows the tether attachment point. **C)** Sequential tether pulling from two points on the same cell (white arrows). Top: pulling from left edge of the cell. The image at t = 0 shows the expression of PIEZO1-mCherry. Bottom: pulling from the right sides of the same cell. Images in **B** and **C** are composites of mean fluorescence (grey) and changes in fluorescence (heat map). Scale bars in all panels 10 μm.

**Fig. S10.**
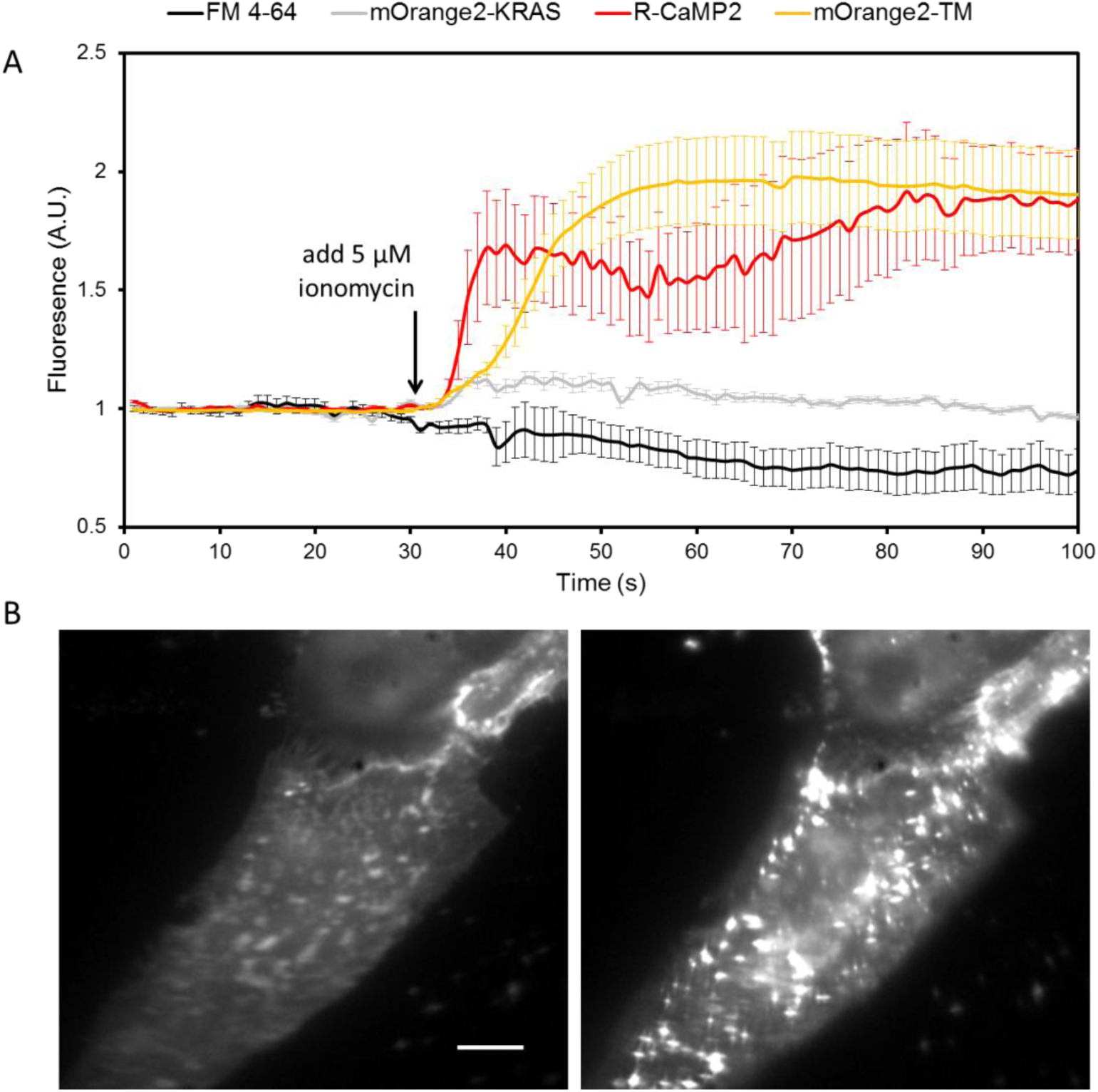
Validating mOrange2-TM as a vesicle fusion reporter. **A)** Validating ionomycin as a means to induce vesicle fusion. In MDCK cells expressing R-CaMP2, ionomycin (5 µM) induced a rapid increase in fluorescence (red, *n* = 4 cells) indicating Ca^2+^ influx. Fresh MDCK cells were incubated with FM 4-64 to load the dye into vesicles and then the dye was washed from the extracellular medium. Ionomycin led to a decrease in fluorescence (black, n = 4 cells), consistent with ionomycin-induced vesicle fusion. In MDCK cells expressing mOrange2-TM, ionomycin induced a rapid increase in fluorescence (orange, n = 7 cells), consistent with vesicle fusion and de-acidification of the vesicles. As a control experiment, ionomycin was added to MDCK cells expressing mOrange2-KRAS, which targeted the fluorescent protein to the inner leaflet of plasma membrane and to the cytoplasmic surface of vesicles. Ionomycin did not affect the fluorescence of these cells (grey, n = 5 cells). Error bars are s.e.m.. **B)** Image of MDCK cells expressing mOrange2-TM before (left) and after (right) adding ionomycin. Scale bar 10 μm.

**Fig. S11.**
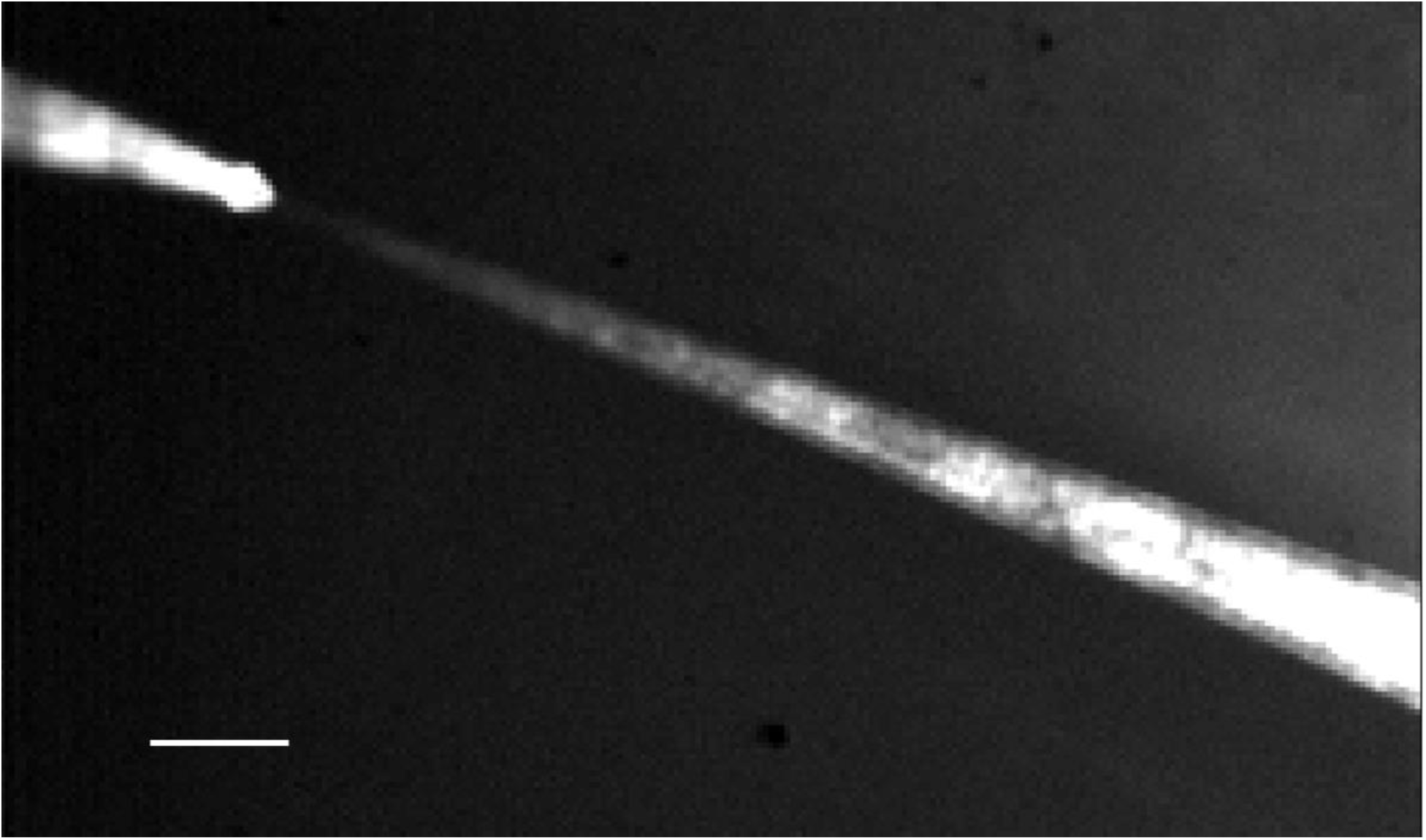
Tracking the flow profile from the pipette for shear perturbation to cells. The pipette is on the upper left. The flow was visualized with fluorescent beads. Scale bar 100 μm.

## Supplementary Table

**Table S1.** Diffusion coefficients of different tracer molecules measured on a tether and a cell body with FRAP as described in Fig. S5. All reported values are mean ± s.e.m. See *Materials and Methods* for details of DNA constructs.

| Tracer molecules | $D_s^{\text{cell}}$ ($\mu\text{m}^2/\text{s}$) | $D_s^{\text{tether}}$ ($\mu\text{m}^2/\text{s}$) | Ratio |
| --- | --- | --- | --- |
| DRD2-eGFP | $0.037 \pm 0.005$ (n=10) | $0.76 \pm 0.08$ (n=10) | $21 \pm 4$ |
| CheRiff-eGFP | $0.035 \pm 0.003$ (n=4) | $0.48 \pm 0.07$ (n=4) | $14 \pm 3$ |
| ASAP1 | $0.024 \pm 0.003$ (n=4) | $0.5 \pm 0.15$ (n=4) | $21 \pm 7$ |
| eGFP-TM | $0.05 \pm 0.01$ (n=3) | $0.81 \pm 0.22$ (n=3) | $16 \pm 5$ |
| GPI-eGFP | $0.10 \pm 0.03$ (n=3) | $1.07 \pm 0.05$ (n=3) | $11 \pm 4$ |

